# Deep Learning-Based High-Resolution Time Inference for Deciphering Dynamic Gene Regulation from Fixed Embryos

**DOI:** 10.1101/2025.03.17.643618

**Authors:** Huihan Bao, Shihe Zhang, Zhiyang Yu, Heng Xu

## Abstract

Embryo development is driven by the spatiotemporal dynamics of complex gene regulatory networks. Uncovering these dynamics requires simultaneous tracking of multiple fluctuating molecular species over time, which exceeds the capabilities of traditional live-imaging approaches. Fixed-embryo imaging offers the necessary sensitivity and capacity but lacks temporal resolution. Here, we present a multi-scale ensemble deep learning approach to precisely infer absolute developmental time with 1-minute resolution from nuclear morphology in fixed *Drosophila* embryo images. Applying this approach to quantitative imaging of fixed wild-type embryos, we resolve the spatiotemporal regulation of the endogenous segmentation gene *Krüppel* (*Kr*) by multiple transcription factors (TFs) during early development without genetic modification. Integrating a time-resolved theoretical model of single-molecule mRNA statistics, we further uncover the unsteady-state bursty kinetics of the endogenous segmentation gene, *hunchback* (*hb*), driven by dynamic TF binding. Our method provides a versatile framework for deciphering complex gene network dynamics in genetically unmodified organisms.

## Introduction

Embryo development is orchestrated by complex gene regulatory networks in individual cells, whose dynamic spatiotemporal expressions dictate cell fate determination and body plan formation^1–3^. For example, in early *Drosophila* embryos (nuclear cycles (nc) 11 to early 14), cell nuclei undergo rapid and synchronous divisions, with no visible body structures yet formed. Yet, during this stage, future body segments along the anterior-posterior (AP) axis are pre-defined by the dynamic expression patterns of the segmentation gene network^4–6^. To unravel these gene regulation dynamics, a key challenge is to simultaneously track both the spatial and temporal information with high resolution. A common approach involves real-time imaging of live embryos genetically engineered to express fluorescently labeled mRNAs or proteins of interest^7–10^. However, due to genetic and optical challenges, this approach inherently struggles to monitor multiple dynamic molecular species simultaneously (typically <4)^11^, limiting its effectiveness in analyzing complex regulatory interactions. Additionally, genetic modifications can disrupt endogenous gene expression, potentially distorting the natural biological system^11^.

Fixed-embryo imaging offers a compelling alternative that does not require genetic modifications and is scalable to high-throughput applications^12–14^. Without the trade-offs associated with live tracking, it provides higher sensitivity and spatial resolution^15^. The challenge, however, is that each fixed-embryo image represents only a snapshot in time, necessitating temporal alignment of multiple fixed embryos to reconstruct the developmental process. Traditionally, this alignment relied on manually timing embryos before fixation^16,17^ or comparing their morphology to a pre-established developmental atlas after fixation^18,19^. Yet, due to uncertainty in developmental start times and the limited sample density of the atlas, these methods provide only coarse-grained developmental stages, inadequate for resolving detailed gene regulation dynamics at minute or sub-minute timescales.

One way to improve temporal resolution is by leveraging fast-changing morphological features at the cellular level^5,20–23^. For example, during the cellularization stage of the *Drosophila* embryo (early-mid nc14), the progressively elongating furrows of invaginating cell membranes have been used to estimate the developmental time with ∼2-4-minute resolution^5,20^. However, in most other cases, the relationship between time and cell morphology is complex and stage-dependent^21–23^. Explicitly identifying these subtle features and reliably mapping them to absolute developmental time remains technically challenging. Deep learning methods, which can objectively extract comprehensive image features and model complex relationships, have recently been applied as a powerful tool to study embryo development^24,25^. However, while most existing applications focus on classifying embryo phenotypes and coarse-grained developmental stages^26–28^, a versatile and high-resolution inference of absolute developmental time is still lacking.

Here, we present a deep learning-based regression approach to infer the absolute developmental time of early *Drosophila* embryos during nc11-early 14 with 1-minute resolution. Using time-lapse nuclear histone images of transgenic embryos as the initial dataset, we employed ensemble learning with three independent convolutional neural network (CNN) models to capture morphological features across multiple spatial scales^29–31^. By calibrating fixation-induced and strain-specific variations in embryo size, and implementing a relayed learning strategy, our method precisely inferred developmental time from standard DNA images of fixed embryos, regardless of strain. Using this approach, we quantified the spatiotemporal regulation of the segmentation gene *Krüppel* (*Kr*) by two transcription factors (TFs), Bicoid (Bcd) and Hunchback (Hb), from single-molecule imaging of multiple fixed wild-type (WT) embryos during nc11-13. We found that *Kr* transcription is dynamically governed by a multiplicative combination of cooperative Bcd activation and Hb repression. Moreover, by incorporating single-molecule mRNA statistics, we developed a time-resolved theoretical pipeline to uncover the microscopic transcription kinetics of the segmentation gene *hunchback* (*hb*) during nc11-13. We found that *hb* transcription follows unsteady-state bursty kinetics driven by dynamic Bcd binding within a specific time window of each nuclear cycle. Our method offers a versatile framework for investigating dynamic gene regulation in genetically unmodified organisms and facilitates spatiotemporal multi-omics for deciphering complex gene regulatory networks.

## Results

### Predicting developmental time from nuclear histone images with 1-minute accuracy

To relate developmental time with dynamic nuclear signal, we obtained time-lapse images of histone H2A-RFP from >30 transgenic *Drosophila* embryos (*his2av-mrfp1*) during nc10-early 14 with 1-minute time-resolution using confocal microscopy (Fig. 1a and Supplementary Figs. 1a-c, see Methods). The timing of different embryos was aligned and scaled based on cell division times to compensate for variation in developmental tempo among embryos. Besides a significant difference in the number of nuclei between nuclear cycles, we observed a continuous evolution of nuclear morphology (including size, shape, etc.) across multiple spatial scales within each nuclear cycle (Fig. 1a and Supplementary Fig. 1d).

**Fig. 1.**
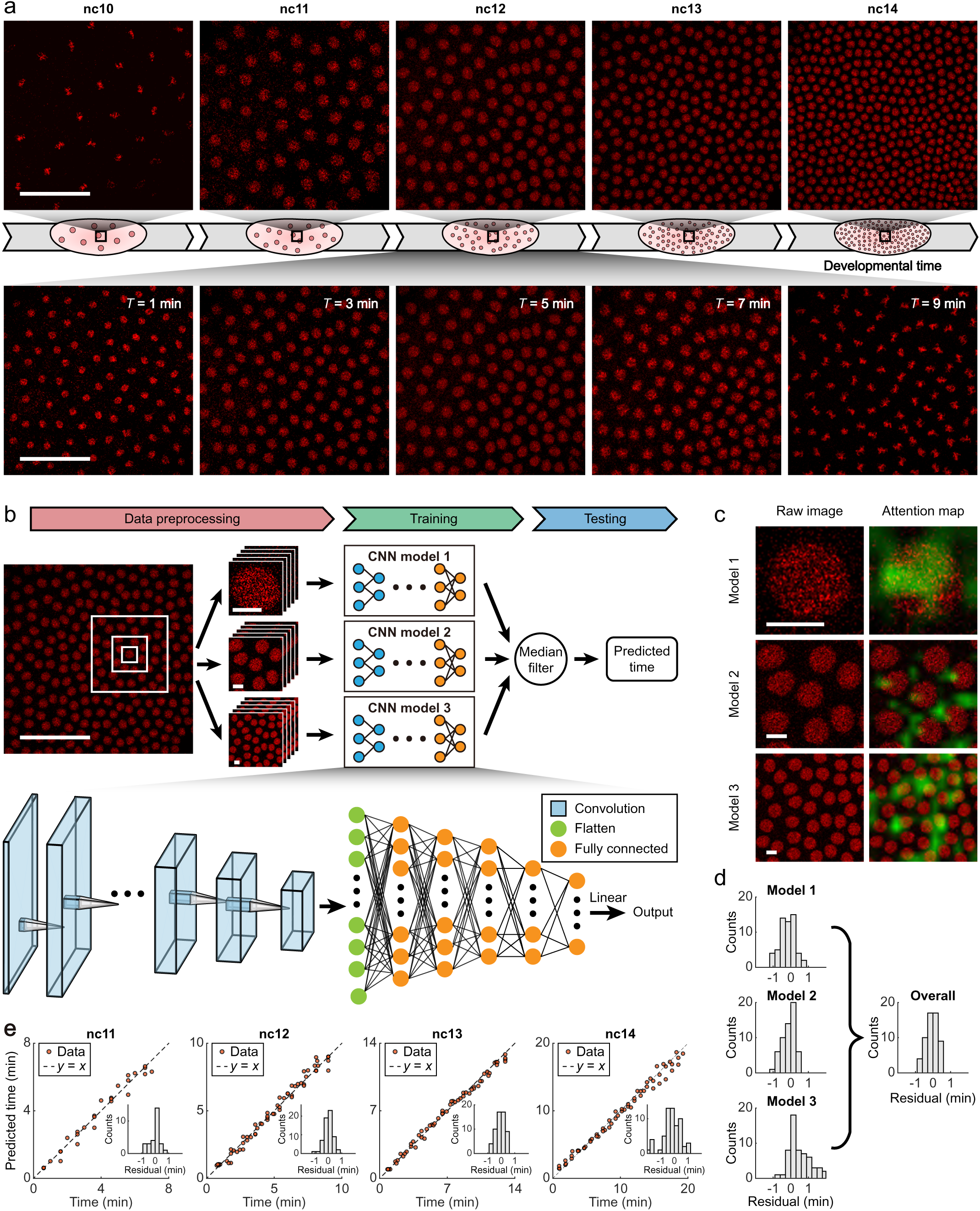
CNN models predict developmental time from nuclear histone images of live embryos. **a,** Still frames from time-lapse images of a *his2av-mrfp1* embryo during nc10-14 (top) and within nc12 (bottom). Scale bar, 50 μm. **b,** Schematics of the time prediction workflow (top) and model architecture (bottom). Scale bars, 50 μm (left) and 5 μm (right). **c,** Grad-CAM visualization of the attention from three independent models. Left: representative histone images from an nc13 embryo at different spatial scales as inputs for each model. Right: Grad-CAM maps (green) from the final convolutional layer in each model, overlaid on the original images. Scale bar, 5 μm. **d,** Histograms of residuals from individual models (left) and the overall time predictor (right) in nc13. **e,** Evaluation of developmental time prediction across multiple nuclear cycles. The predicted time for each time point of each embryo was compared to ground truth. Insets: histograms of residuals from the overall time predictor. *n* = 4, 6, 4, and 3 embryos for nc11-14, respectively.

To comprehensively extract all time-dependent features from embryo images, we constructed a multi-scale ensemble deep learning framework consisting of three independent VGG-like CNN models, each with a regression output layer for continuous-time prediction^30,31^ (Fig. 1b, Supplementary Table 1, see Methods). By dividing every embryo image into multiple small windows of three different sizes covering single, few (∼6), and multiple (∼20) nuclei, respectively, we trained each CNN model independently to capture time-dependent features at different spatial scales (Fig. 1c and Supplementary Fig. 1e). At any time point during nc11-early 14, predictions from these models were always similar and closely matched the ground truth, mutually validating each other (Supplementary Fig. 1f). Further combining the three models through a median filter outperformed any individual predictions (Fig. 1d and Supplementary Fig. 1f).

Compared with ground truth, our time prediction is both accurate and precise across all nuclear cycles, with little bias and variability (mean residual ± SD, nc11: 0.01 ± 0.35 min, nc12: 0.02 ± 0.34 min, nc13: 0.05 ± 0.34 min, nc14: −0.2 ± 0.68 min, Fig. 1e). Notably, almost all predictions for nc11-early 14 are within 1-minute accuracy (nc11: 100%, nc12: 98%, nc13: 100%, nc14: 87%), catching the imaging time resolution. In particular, early nc14 predictions significantly outperformed the traditional time inference method based on membrane invagination^5,20^ (∼2-4-minute accuracy), demonstrating the power of our deep learning method. Given this enhanced accuracy, we focus on inference for nc11-13 in the following study, as no other methods exist for this period.

### Size rescaling enables time inference from fixed-embryo imaging

To apply the trained time inference models to fixed embryos, we compared images from fixed and live embryos of the same strain (*his2av-mrfp1*) during nc11-13. Consistent with previous reports of fixation-induced embryo shrinkage^20,32^, we observed a decrease in nuclear size of a similar proportion (∼80%-90%, Fig. 2a). Since nuclear size is one of the major time-dependent features captured by our models, such shrinkage severely disrupted time inference accuracy (Supplementary Figs. 2a-b).

**Fig. 2.**
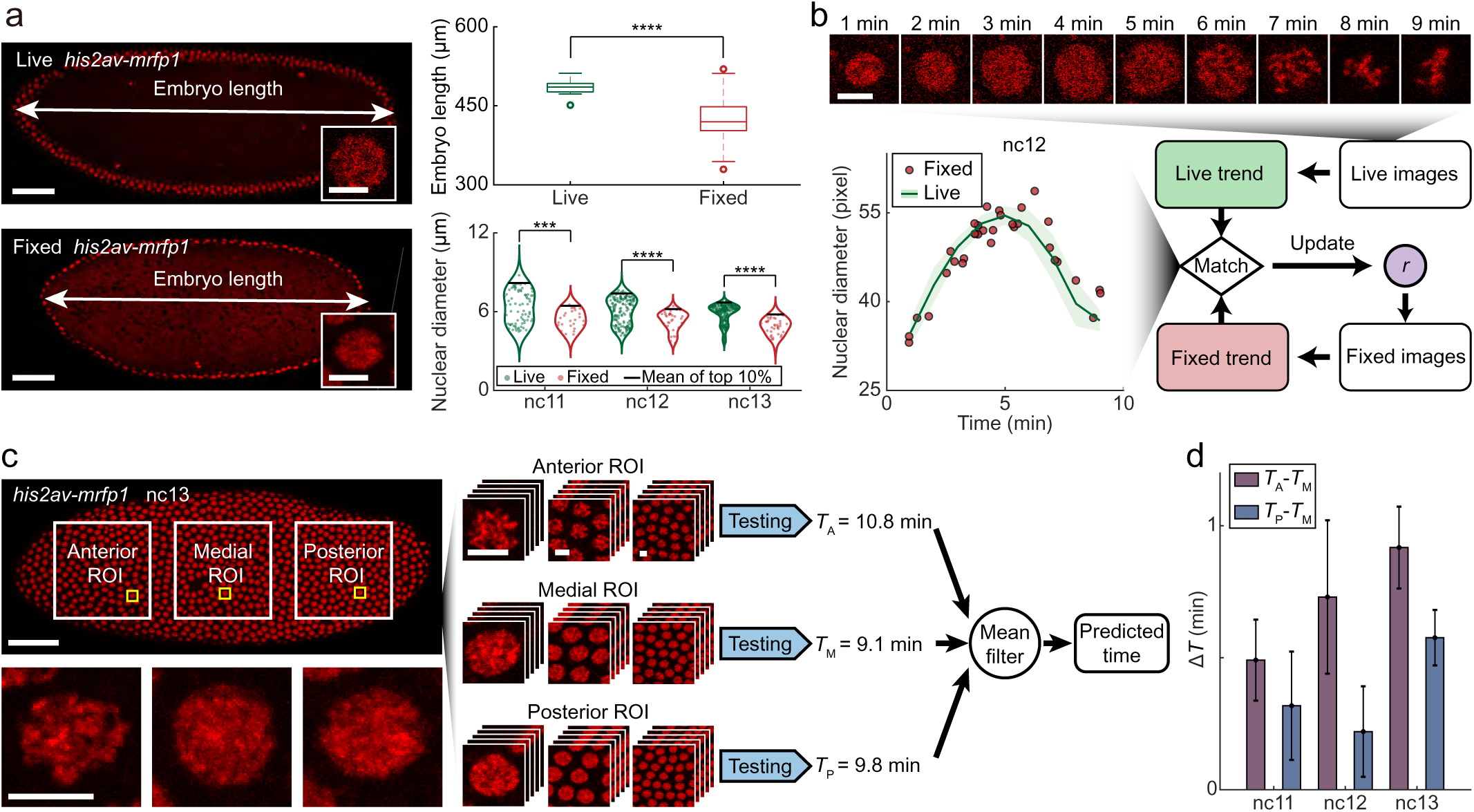
Time inference from histone images of fixed embryos with image size rescaling. **a,** Fixation induces embryo shrinkage. Left: confocal images of live (top) and fixed (bottom) *his2av-mrfp1* embryos at the mid-sagittal plane. Scale bars, 50 μm. Insets, magnified views of individual nuclei. Scale bars, 5 μm. Top right: size comparison between live (*n* = 14) and fixed (*n* = 155) embryos, with two-sided *t*-test (\*\*\*\**p* < 0.0001). Bottom right: comparison of nuclear diameter between live (*n* = 11, 18, and 22 embryos for nc11-13) and fixed embryos (*n* = 28, 33, and 38 embryos for nc11-13), with two-sided *t*-test (\*\*\**p* < 0.001; \*\*\*\**p* < 0.0001). **b,** Schematic of determining the optimal rescaling magnitude. Live histone images over nc12 (top) were used to extract the trend of nuclear diameter over time (bottom left, green line, data from *n* = 18 live embryos). This trend was compared iteratively with reconstructed data from rescaled fixed-embryo images (bottom left, red dots, data from *n* = 33 fixed embryos) to optimize the rescaling magnitude. Scale bar, 5 μm. Shading indicates SD. **c,** Schematic of independent time inferences for anterior, medial, and posterior regions of a fixed *his2av-mrfp1* embryo at nc13. Magnified views of representative nuclei from each region are shown below. Right: results from different regions were averaged for an overall time prediction. Scale bars, 50 μm (embryo) and 5 μm (magnified views). **d,** Differences in inferred times between anterior and medial regions, and between posterior and medial regions, during late mitotic interphase for each nuclear cycle (mean ± s.e.m.). Anterior-medial difference: *n* = 10, 5, and 10 embryos for nc11-13, respectively; posterior-medial difference: *n* = 7, 10, and 6 embryos for nc11-13, respectively.

To correct this effect, we preceded our time inference framework with an image-rescaling step for fixed embryos (see Methods). To determine the magnitude of rescaling, we used the trend of nuclear size over time as a reference (Fig. 2b). For each nuclear cycle, by directly measuring this trend from live imaging, we tried to reconstruct it from a set of fixed embryos through time inference with various rescaling magnitude (Fig. 2b, Supplementary Figs. 2c-d, see Methods). The best reconstructions matched well with live measurement results (Supplementary Fig. 2d), indicating an optimal rescaling magnitude of ∼1.20, consistent with previous reports^32^.

By applying this time inference to individual nuclei in different regions of fixed embryos at mitotic interphase, we observed temporal asynchrony along the AP axis (Fig. 2c). This is likely a residue of the mitotic wave, where nuclear divisions occur in a wave-like pattern that moves from the poles to the equator of the *Drosophila* embryo within ∼0.5-1 min^33,34^. Specifically for embryos in late mitotic interphase of nc11-12, the inferred time for the medial region (∼0.5 embryo length (EL)) exhibited a significant delay of ∼0.3-0.6 min compared to the anterior and posterior regions (∼0.2 EL and ∼0.8 EL). In nc13, this time delay increased to ∼0.6-1.0 min, consistent with previous reports of wave slow-down during this cycle^33,34^ (Fig. 2d, Supplementary Fig. 2e). These results demonstrate the 1-minute resolution of our time inference method for fixed embryos. In subsequent applications, to mitigate the influence of the mitotic wave, we averaged the inference results from these three regions to generate an overall time prediction for each fixed embryo (Fig. 2c).

### Relayed learning enables time inference from the nuclear DNA signal of WT embryos

Beyond histone, which requires genetic modification of fly strains or specific antibodies for labeling, nuclear DNA, which can be easily marked by organic dyes^35,36^, is a rather common target for fixed-embryo imaging. However, directly training our framework for the DNA signal is challenging due to the difficulty of labeling and imaging nuclear DNA in live embryos. We thus used the histone signal in fixed embryos as a relay to link the nuclear DNA signal with developmental time for training (see Methods). Specifically, by simultaneously imaging histone and DNA signals from 160 fixed *his2av-mrfp1* embryos during nc11-13 (Fig. 3a), we inferred embryonic times from histone signals and used them as input to train and test three DNA-based models at different spatial scales (Fig. 3b). With proper calibration of image contrast and saturation (Supplementary Figs. 3a-c), the combined predictions from these DNA-based models closely matched histone-based time inference, achieving 1-minute accuracy and high precision in all cycles (mean residual ± SD, nc11: 0.20 ± 0.46 min, nc12: 0.14 ± 0.47 min, nc13: 0.19 ± 0.42 min, Fig. 3c and Supplementary Fig. 3d).

**Fig. 3.**
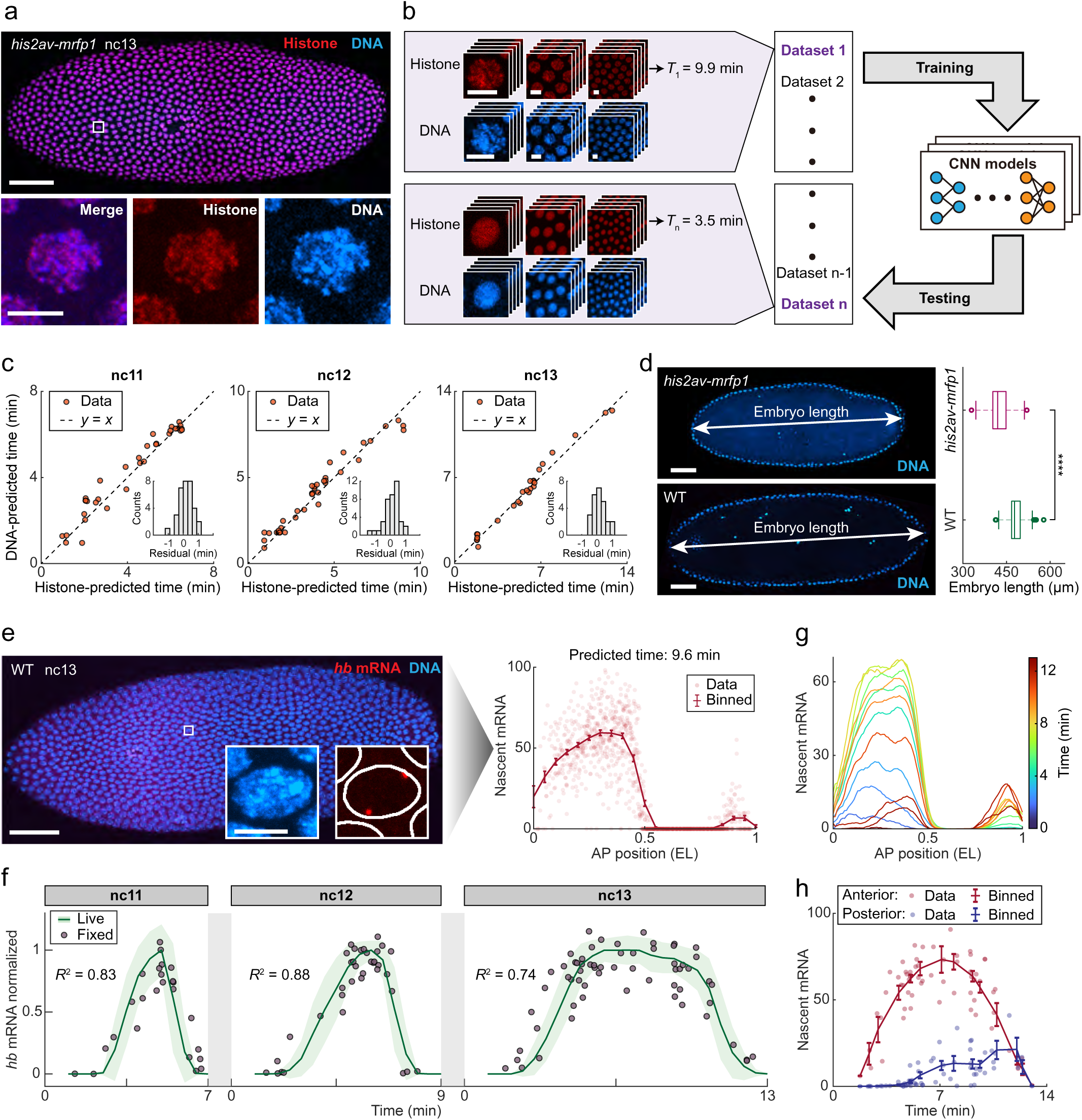
Time inference from DNA images of fixed embryos using relayed learning. **a,** Confocal image of a fixed *his2av-mrfp1* embryo labeled for DNA at nc12, with magnified views of a single nucleus displayed below. Scale bars, 50 μm (embryo) and 5 μm (magnified views) **b,** Workflow of relayed learning. Developmental time inferred from histone images was used to train and test DNA-based models. Scale bar, 5 μm. **c,** Evaluation of DNA-based time prediction across multiple nuclear cycles. DNA-predicted times for multiple fixed embryos were compared to histone-predicted times. Insets: histograms of residuals from the overall time predictor. *N* = 33, 40, and 25 embryos for nc11-13, respectively. **d,** Embryo size difference between strains. Left: confocal images of fixed *his2av-mrfp1* and WT embryos at the mid-sagittal plane. Scale bar, 50 μm. Right: size comparison between fixed *his2av-mrfp1* (*n* = 155) and WT (*n* = 100) embryos, with two-sided *t*-test (\*\*\*\**p* < 0.0001). **e,** Confocal image of a fixed WT embryo labeled for hb mRNA and DNA at nc13. Scale bar, 50 μm. Insets: magnified views of a single nucleus. Scale bar, 5 μm. Right: number of nascent *hb* mRNAs per nucleus as a function of the AP position. Data were binned along the AP axis (bin size: 0.1 EL, step size: 0.05 EL) to show mean ± SD. **f,** Comparison of *hb* transcription dynamics (averaged over 0.2-0.4 EL) extracted from live (*his2av-mrfp1*) and fixed (WT) embryos across nc11-13. Live-imaging data from individual embryos were max-normalized, averaged across multiple embryos (*n* = 4, 4, and 9 embryos for nc11-13, respectively) along the time axis (bin size: 1 min, step size: 0.5 min), and then max-normalized again for each nuclear cycle. Shadings indicate SD. Fixed-imaging data (*n* = 25, 36, and 63 embryos for nc11-13, respectively) were rescaled to match live *hb* trend for each nuclear cycle. **g,** Spatial profile of endogenous *hb* transcription over time during nc13, binned along the AP-axis (bin size: 0.06 EL, step size: 0.01 EL) and time (bin size: 2 min, step size: 1min). **h,** Maximum transcription levels of anterior (0-0.5 EL) and posterior (0.8-1 EL) *hb* expression bands over time during nc13. Data from individual embryos were time-binned (bin size: 2 min, step size: 1min) to show mean ± s.e.m. **g-h,** Data from 63 fixed WT embryos.

To generalize this DNA-based time inference for other fly strains, particularly the WT, we noticed systematic differences in embryo size between different strains (Fig. 3d)^23,32^. Thus, DNA images of WT embryos need to be rescaled accordingly before applying the *his2av-mrfp1*-trained framework (see Methods). To validate this approach, we simultaneously imaged nuclear DNA and nascent mRNA of the anteriorly expressed *hb* gene in 124 fixed WT embryos during nc11-13 (Fig. 3e, see Methods). By inferring the developmental time and quantifying the anterior *hb* mRNA level (0.2-0.4 EL) for each embryo, we reconstructed the endogenous *hb* transcription dynamics from fixed WT embryos without requiring genetic modification (Fig. 3f, Supplementary Figs. 3e-f). This dynamics quantitatively matched the live imaging results of *hb-*MS2 in transgenic embryos (see Methods), demonstrating the accuracy and universal applicability of our DNA-based framework in different fly strains.

Further analyzing *hb* transcription along the entire AP axis obtained a complete spatiotemporal profile of endogenous *hb* transcription in WT embryos (Fig. 3g, Supplementary Fig. 3g, see Methods). This is a critical complement to previous live imaging studies, which had limited field-of-view and often omitted the posterior *hb* expression band^7,37^. Here, we observed that the posterior expression band emerged later than the anterior one, with significantly delayed initiation and peak times (Fig. 3h). This finding aligns with previous reports that anterior *hb* transcription is activated by early-produced maternal Bcd, while posterior *hb* transcription is regulated by zygotic transcription factors, e.g., Tll^38^, which are produced much later^4,39^ (Supplementary Fig. 3h). In contrast, both bands vanished simultaneously, suggesting a shared mechanism of transcription termination.

### Time inference of fixed embryos resolves the dynamic regulation of *Kr* by multiple TFs

Transcriptional regulation involves complex and dynamic interactions between multiple factors over time. Our time inference method, combined with multiplex fluorescence imaging of mRNA and regulatory factors in fixed embryos, enables direct measurement of such regulatory dynamics. Here, we used a multi-factor-regulated gene, *Kr*, as an example to demonstrate this approach. According to previous studies, the transcriptional regulation of *Kr* in early development (nc11-13) primarily involves two maternal TFs, i.e., an activator, Bcd, and a repressor, Hb, whose regulatory dynamics remained unknown^4,40,41^. We thus applied smFISH^35,36,42,43^, immunofluorescence^5,20,42^, and a DNA dye to simultaneously label *Kr* mRNA, Bcd and Hb proteins, and nuclear DNA in >130 fixed WT embryos during nc11-13 (Fig. 4a, see Methods).

**Fig. 4.**
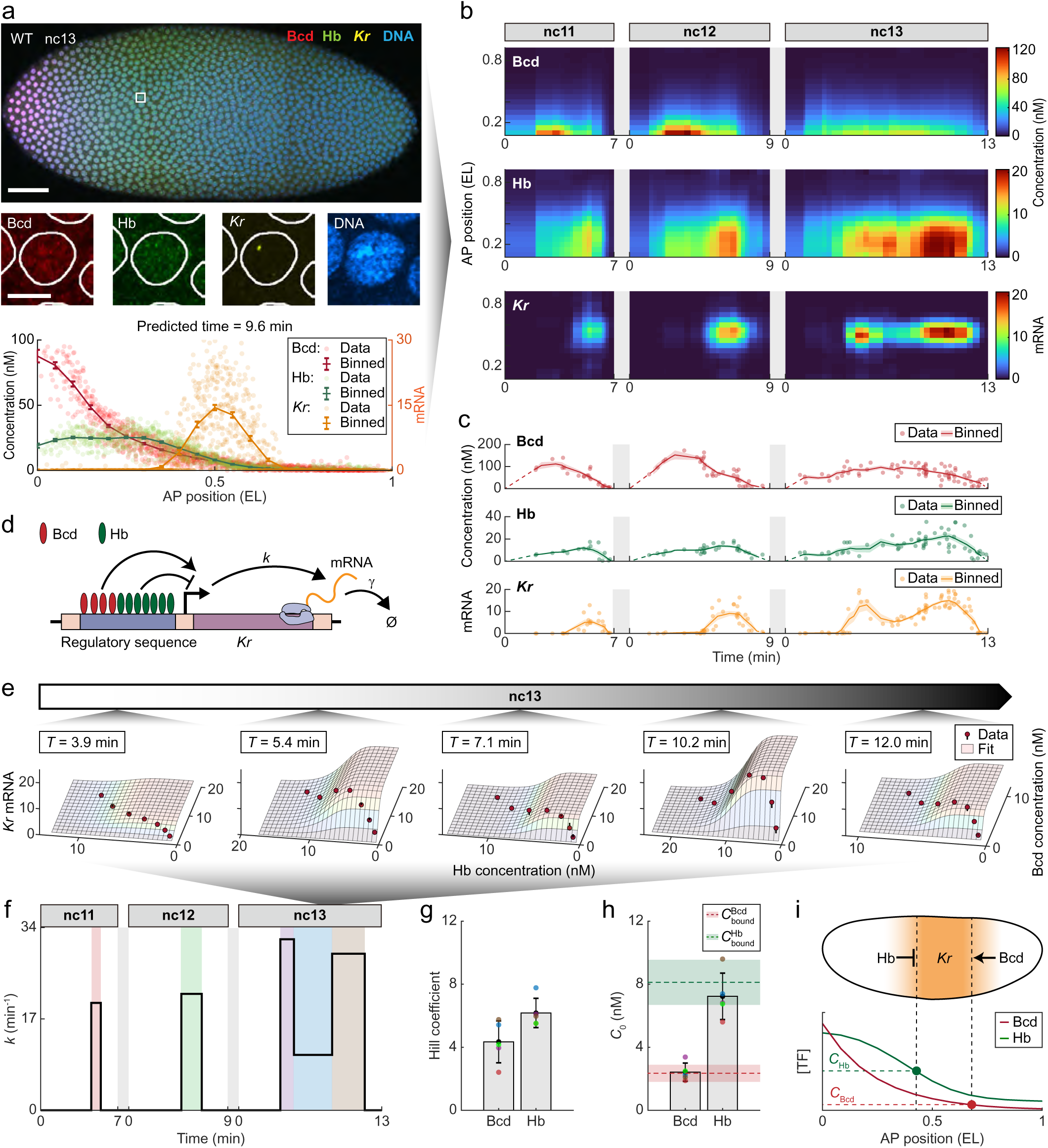
Resolving the dynamic regulation of *Kr* by multiple TFs from fixed embryos. **a,** Confocal image of a fixed WT embryo labeled for Bcd protein, Hb protein, *Kr* mRNA and DNA at nc13, with magnified views of a single nucleus displayed below. Scale bars, 50 μm (embryo) and 5 μm (magnified views). Bottom: Bcd and Hb concentrations and number of nascent *Kr* mRNAs per nucleus as a function of the AP position. Data were binned along the AP axis (bin size: 0.1 EL, step size: 0.05 EL) to show mean ± SD. **b,** Spatiotemporal profiles of nuclear Bcd concentration, Hb concentration, and *Kr* transcription in nc11-13. Data were binned along the AP axis (bin size: 0.1 EL, step size: 0.05 EL) and time (bin size: 1 min, step size: 0.25, 0.25, and 0.5 EL for nc11-13, respectively). **c,** Maximum Bcd and Hb concentrations along the AP axis and the average transcription level of *Kr* expression band (0.4-0.6 EL) as functions of time during nc11-13. Data from individual embryos were binned along the time axis (bin size: 2 min, step size: 0.5 min). Shadings indicate s.e.m. **d,** Schematic of endogenous *Kr* regulation through cooperative Bcd and Hb binding at enhancers, with Bcd functioning as an activator and Hb as a repressor. **e,** Regulatory relationship between *Kr*, Bcd, and Hb over time in nc13. AP axis-binned data from (**b**) (red dots) were fitted to a time-dependent thermodynamic model (surfaces). **b-c, e,** Data from *n* = 25, 36, and 76 fixed WT embryos for nc11-13, respectively. **f,** mRNA production rate as a function of time. Five expression periods (*k* > 0) in nc11-13 were shown in distinct colors. **g-h,** Comparison of Hill coefficients (**g**) and concentration thresholds (**h**) for Bcd and Hb. Values from different expression periods (dots) were summarized as mean ± SD. Bcd and Hb concentrations at average *Kr* boundaries for WT embryos (*n* = 68) during the activatable periods (*k* > 0) in nc11-13 (mean ± SD) are shown as reference. **i,** Schematic showing the determination of anterior and posterior *Kr* expression boundaries by the spatial profiles of Hb and Bcd, respectively.

By inferring developmental time from DNA signals, we quantified the spatiotemporal profiles of nuclear *Kr* transcription and Bcd and Hb concentrations in individual embryos over nc11-13 (Fig. 4b and Supplementary Figs. 4a-c, see Methods). Within each mitotic interphase, *Kr* formed a medial expression stripe (∼0.45-0.60 EL), while Bcd and Hb displayed exponential and reverse sigmoidal gradients along the AP axis, respectively (Figs. 4a-b and Supplementary Fig. 4d), consistent with previous reports^4^. Temporally, all three species varied significantly within each nuclear cycle, with Bcd and Hb levels rising ahead of *Kr* (Fig. 4c). Notably, during nc13, *Kr* exhibited two distinct expression peaks that were previously indistinguishable^41^, highlighting the sensitivity of our method. The duration of the earlier peak was significantly shorter the later one, explaining the previously reported increase in *Kr* expression during late nc13^41^.

To understand the dynamic regulation of *Kr* by Bcd and Hb, we further analyzed the quantitative relationship between the three profiles. Given the presence of multiple Bcd and Hb binding sites on *Kr* enhancers (Supplementary Fig. 4e)^40,44,45^, we constructed a time-dependent thermodynamic model to describe the nuclear mRNA level of *Kr* (*R*) in response to Bcd and Hb concentrations (*C*_Bcd_ and *C*_Hb_) (Fig. 4d, see Methods). Applying the model to fit the spatiotemporal relationship between *Kr*, Bcd, and Hb over nc11-13 yielded (Fig. 4e)

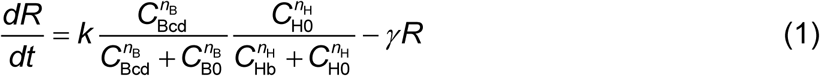

where *n*_B_ and *n*_H_ are Hill coefficients describing the cooperativities of Bcd and Hb bindings, respectively; *C*_B0_ and *C*_H0_ are concentration thresholds for Bcd and Hb regulations, respectively; *k* and *γ* represent mRNA production and degradation rates, respectively (see Methods). This result suggests a synergistic regulatory mechanism, where cooperative Bcd and Hb bindings combine their regulatory effects multiplicatively.

From the fitting, we found that *Kr* was only active (*k* > 0) in a fraction of each nuclear cycle (Fig. 4f). Notably, in nc13, *k* exhibited two pulses, suggesting gene activation in G1 and G2 phases, respectively. This phenomenon was not observed in nc11-12, possibly due to the transiency of G1 in early cycles. Throughout all cycles, *n*_B_ and *n*_H_ remained stable at ∼4 and ∼6, respectively (Fig. 4g), revealing higher-order cooperativity of Hb than Bcd, consistent with bioinformatic studies (Supplementary Fig. 4e)^40,44,45^. *C*_B0_ and *C*_H0_ were estimated to be ∼2.4 nM and ∼7.2 nM, respectively (Fig. 4h), suggesting that Hb repression and Bcd activation determine the anterior and posterior boundaries of *Kr* expression, respectively (Fig. 4i). Further measuring *Kr* regulation in transgenic embryos with reduced Bcd and Hb dosage (1×*bcd* strain) confirmed these findings (Supplementary Fig. 4f-i), revealing that maternal Bcd and Hb are sufficient for early *Kr* patterning.

### Time-resolved single-molecule mRNA statistics reveal unsteady-state kinetics of *hb* transcription

Beyond population-level studies, fixed-embryo imaging can achieve single-molecule sensitivity, enabling statistical analysis to uncover the microscopic mechanisms of gene regulation. Following this strategy, previous smFISH studies of nascent mRNA copy-number distribution at individual gene loci have identified the bursty nature of stochastic gene transcription in various biological systems^35,46,47^. However, most of these studies assumed gene activity at steady state^35,46,47^, which is inadequate for highly dynamic systems, e.g., developing embryos. Here, with accurate time inference, we used the *hb* gene as an example to establish a time-resolved theoretical pipeline for deciphering unsteady-state transcription kinetics from multiple smFISH-labeled embryos covering nc11-13 (Fig. 5a, see Supplementary Note 1).

**Fig. 5.**
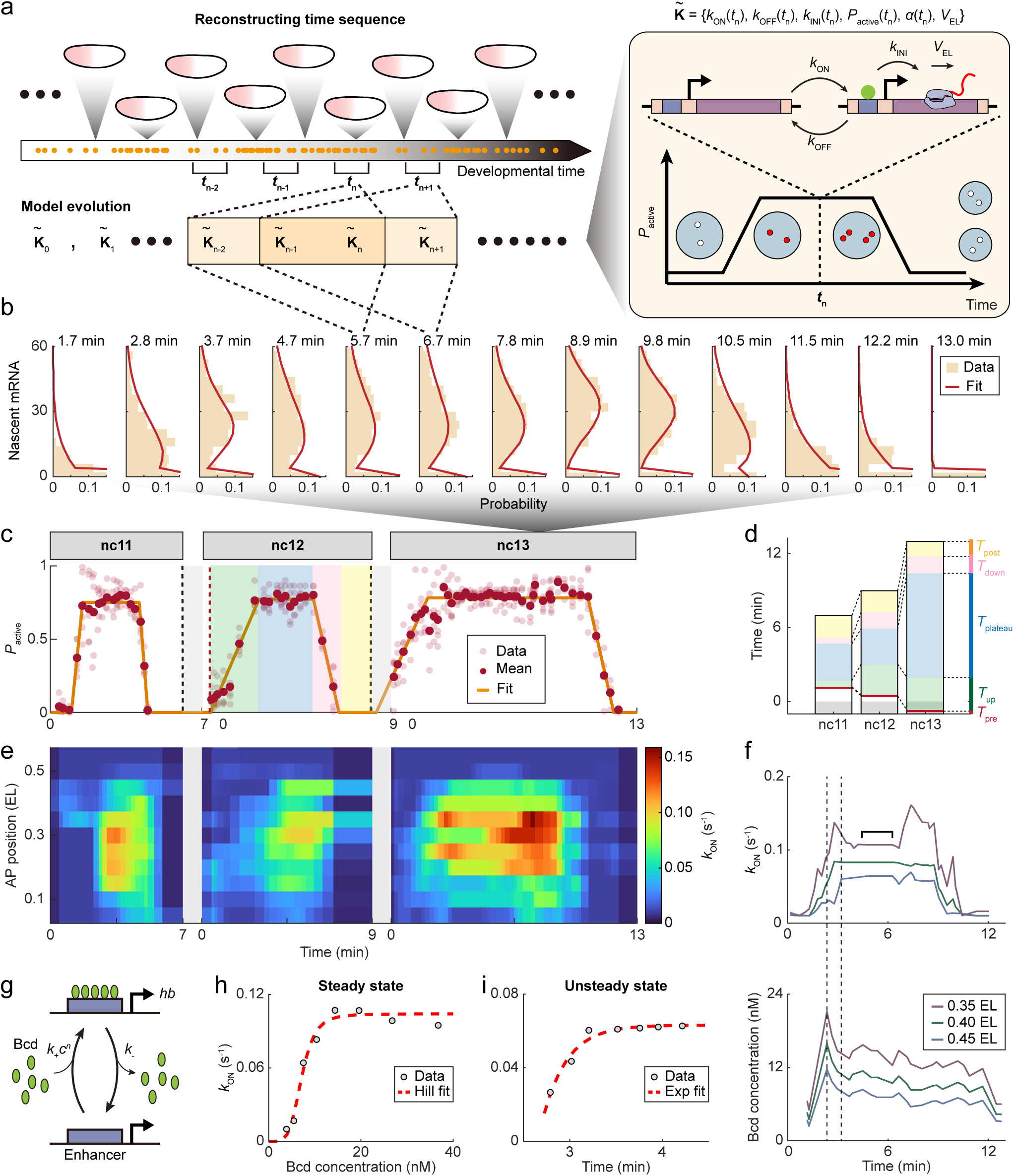
Uncovering *hb* transcription kinetics through time-resolved single-molecule mRNA statistics. **a,** Schematic of the theoretical pipeline for deciphering unsteady-state transcription kinetics. Time inference for multiple smFISH-labeled fixed embryos enables reconstructing the evolution of single-cell mRNA signals (left), which were fitted to a time-dependent two-state transcription model (right). The mRNA signal at a time point is influenced by kinetic parameters within a preceding time window. **b,** Histograms of nascent *hb* mRNAs at individual anterior gene loci (0.25-0.35 EL) over time in nc13. Data from *n* = 63 WT embryos were binned along the time axis (bin size: 0.5 min, step size: 0.5 min). All histograms were simultaneously fitted to the time-dependent two-state transcription model. **c,** Percentage of activatable gene loci as a function of time during nc11-13. Data from various AP positions (0.2-0.4 EL) were averaged and fitted with a trapezoidal curve. The rising, plateau, falling, and silent phases of the trapezoidal curve for nc12 are highlighted in different colors. **d,** Comparison of each trapezoidal phase across nc11-13. **e,** Spatiotemporal profile of gene activation rate across nc11-13, for the AP position range of 0.05-0.55 EL. **f,** Promoter activation rate and nuclear Bcd concentration at three distinct AP positions as functions of time in nc13. Dashed lines mark the period during which the Bcd concentration reaches its maximum, while the promoter activation rate continues to rise. Horizontal square bracket marks the period for subsequent steady-state analysis. **g,** Schematic of cooperative Bcd binding-induced gene activation. **h,** Steady-state promoter activation rate at different AP positions (0.20-0.55 EL) during 4.4-6.3 min in nc13 was plotted against nuclear Bcd concentration and fitted to a Hill function. **i,** Unsteady-state promoter activation rate within the marked time period in (**f**) (2.8-4.2 min, 0.45 EL) was plotted against time and fitted to an exponential function.

Based on previous steady-state analyses, *hb* transcription satisfies two-state stochastic kinetics^36,42,48^, with random transitions between active and inactive gene states occuring at Poissonian rates *k*_ON_ and *k*_OFF_. Nascent mRNA molecules are only initiated in the active state at a rate *k*_INI_, followed by elongation at a speed *V*_EL_ and a rapid release. Cooperative Bcd binding activates anterior *hb* transcription by modulating *k*_ON_. To extend this picture to unsteady states, we treated the kinetic parameters of the two-state model as time-dependent variables (Fig. 5a, see Methods). Particularly, to describe the shutdown and re-expression of *hb* during each mitosis (Fig. 3f), we assumed that the percentage of activatable gene loci, *P*_active_, varies with nuclear cycle phases^49^. Notably, since each nascent mRNA signal reflects transcription events within several minutes, signal statistics at successive time points are correlated through inherited kinetic parameters. Retaining this correlation requires simultaneous analysis of the entire time sequence.

Following this extension, we solved the unsteady-state distribution of nascent mRNA number per gene loci over time in different parts of the embryo and compared it with experimental data to extract kinetic parameters (Fig. 5b, see Supplementary Note 1). Unlike steady-state analyses that treated *V*_EL_ as a predetermined scaling factor^35,36,42,48^, our approach allowed a direct estimation of *V*_EL_ at ∼40 bp/s (Supplementary Fig. 5a), aligning with live-imaging measurements^50,51^. In each nuclear cycle, we found a consistent trapezoidal *P*_active_ profile across the anterior *hb* expression domain (Fig. 5c), agreeing with live-imaging results^49^. The onset of this trapezoidal profile advanced with the nuclear cycle, while its duration increased markedly, indicating a rise in gene activity throughout development (Fig. 5d).

During each nuclear cycle, we found that *k*_ON_ varied significantly with AP position and time (Fig. 5e), while *k*_OFF_ and *k*_INI_ remained stable (Supplementary Figs. 5b-c). These results corroborate previous steady-state analyses, confirming that *k*_ON_ is the primary target of Bcd modulation^35,42^. Specifically, at each AP position, the temporal profiles of *k*_ON_ and Bcd concentration both showed rising, plateau, and falling phases (Fig. 5f). During the plateau phase, the steady-state Bcd dependence of *k*_ON_ followed a Hill function with a Hill coefficient of ∼5, consistent with the canonical picture of cooperative Bcd binding-induced gene activation^42^ (Figs. 5g-h). In contrast, the rapid rising phase exhibited unsteady-state behaviors. Notably, we observed a continued increase in *k*_ON_ for ∼1.5 min after the Bcd concentration peaked (Fig. 5f). This relaxation behavior can be explained by extending the canonical model to unsteady state (Fig. 5i, see Supplementary Note 1), which allowed a direct estimation of Bcd binding and unbinding rates (see Supplementary Note 1). During the falling phase, *k*_ON_ decreased ahead of Bcd (Fig. 5f), suggesting a TF-independent mechanism of transcription termination, similar to *Kr* (Fig. 3h). Collectively, these results demonstrate the power of our method in elucidating kinetic mechanisms of gene regulation.

## Discussion

Direct spatiotemporal tracking of complex developmental gene regulation through live imaging has long been limited to monitoring only a few factors simultaneously^11^. Although recent advances in fluorescence lifetime imaging offer potential solutions^52^, the genetic perturbations from multiple fluorescent reporters complicate this approach. Conversely, fixed-embryo imaging could overcome these limitations but lacks precise temporal resolution. In this paper, we present a CNN-based deep learning approach to objectively infer the absolute development time from the nuclear signals in fixed-embryo images with 1-minute resolution. Using early *Drosophila* embryos as a case study, we demonstrate its power in unraveling transcriptional regulation of key developmental genes at the single-cell and single-molecule level.

Compared to existing CNN-based embryo staging, our method offers several key improvements: (1) Instead of analyzing macroscopic embryo morphology that varies slowly, we focus on fast-changing nuclear signals, enabling finer temporal resolution. (2) Unlike traditional CNN-based staging that produces classification outputs^26,27^, our method uses regression outputs, allowing for continuous time prediction. (3) Rather than using a single neural network, whose learning may be limited to a specific spatial scale^26,27^, we employ three independent networks to cover multiple spatial scale. Compared to other multi-scale approaches, such as feature fusion^53,54^, our combination of three independent predictions serves as an ensemble validation, enhancing robustness and providing greater scalability for future adaptation to other systems. (4) To apply our method to fixed embryos of arbitrary strains, we rescale images to compensate for fixation-induced and strain-specific variations in nuclear size –a key time-dependent feature captured by our models. (5) To extend the applicability of our method to DNA images, we use fixed embryos colabeled with histone and DNA as intermediates to train DNA-based models. Overall, these improvements ensure precise and reliable time inference for fixed embryos of any strain.

Besides morphology-based embryo staging, gene expression and epigenetic features have also been used for staging^6,17^. However, these approaches typically require transcriptomic and/or epigenomic data collected at multiple time points, incompatible with regular imaging studies. Moreover, because gene expression is highly influenced by genetic modifications, the versatility and reliability of these approaches across different strains are uncertain. Conversely, our morphology-based method is more robust against genetic manipulation.

Previous fixed-imaging studies of gene regulation are typically limited to resolving static or quasi-static regulatory relationships^6,35,41,42^. Our method enables, for the first time, high-resolution reconstruction of dynamic gene regulation from fixed WT embryos, capturing dynamic details comparable to live imaging. Without genetic modifications and maturation delays of fluorescent proteins, it offers more accurate quantification^11,55,56^. This allowed us to resolve the dynamic regulatory relationship between *Kr*, Bcd, and Hb, extending the static thermodynamic model of transcriptional regulation^57^. Our results quantitatively verify a previously uncertain hypothesis of combinatorial *Kr* regulation: Bcd and Hb each perform cooperative binding independently, while their regulatory effects combine synergistically through multiplication^58^. These findings underscore the potential of our approach to unravel complex gene regulatory mechanisms.

Single-molecule statistical analysis of fixed-imaging data can uncover microscopic mechanisms of gene regulation^35,46,47^. However, the lack of temporal resolution hinders its application to unsteady-state process^59,60^. With precise time inference, we extended this approach to reconstruct the time-dependent transcription kinetics of individual *hb* gene loci in WT embryos. Our analysis reveals the variation in the percentage of activatable gene loci within the nuclear cycle, a feature previously reported only in live imaging^49^. Comparing the extracted *hb* kinetics with the Bcd profile further extended our understanding of TF binding-induced gene activation to realistic unsteady-state scenario. These results highlight the power of our approach to unravel the microscopic kinetics of dynamic gene regulation.

Although our method requires imaging a large number of embryos, it offers an effective and universal solution for accurately quantifying developmental dynamics without genetic modifications. By imaging other features and retraining the models, it can be easily adapted to other developmental stages, other organisms, or even non-developmental processes. Future integration of this method with high-throughput imaging of RNAs, proteins, and gene loci will further enhance our understanding of complex gene regulatory networks^12–14^.

## Methods

### Fly strains

Oregon-R (OreR) strain was used as the wild type. Histone H2A-RFP strain (*his2av-mrfp1*) was obtained from the Bloomington Drosophila Stock Center (stock number 23650, 23651). Two strains for live imaging of *hb* activity (*yw; histone-rfp; mcp-nonls-gfp* and *yw; hb BAC>ms2*) were developed previously^8^ and were obtained as gifts from Dr. Hernan H. Garcia (University of California at Berkeley). 1×*bcd* strain (+/*CyO-bcd+*; *E1s*) was developed previously^61^ and was obtained as a gift from Dr. Jun Ma (Zhejiang University).

### smFISH Probe design

Sets of DNA oligonucleotides complementary to the target transcripts (48 probes for *hb*, 33 probes for *Kr*) were designed and synthesized as previously reported^35^. *Kr* probes were conjugated with tetramethylrhodamine (TAMRA; Thermo Fisher Scientific, C6123). *hb* probes were conjugated with either Alexa Fluor™ 647 (Invitrogen, A20106) or TAMRA.

### Live imaging sample preparation and data acquisition

*his2av-mrfp1* embryos were collected directly. Female virgins from line *yw; histone-rfp; mcp-nonls-gfp* were crossed with males from line *yw; hb BAC>ms2* for embryo collection. Embryos collected at 25 °C were dechorionated using bleach, mounted between a semipermeable membrane (Biofolie; In Vitro Systems & Services) and a coverslip, and embedded in Halocarbon 27 oil (Sigma), following previous literature^7^.

Live embryos were imaged using a Zeiss LSM 710 and 880 confocal microscopes equipped with a 63×/1.4 NA oil immersion objective in confocal or fast airyscan mode. For *his2av-mrfp1* embryos, 16-bit image sequences were acquired at 1 AU pinhole aperture with a pixel size of 132 nm and a *z*-step size of 1 μm. Most confocal-mode images consisted of 1024 × 1024 × 15 pixels. An exception of five image sequences was captured at 512 × 512 × 15 pixels. Airyscan-mode images were captured at 1012 × 1012 × 19 pixels. The standard temporal resolution was one frame per minute (fpm). An exception of a short image sequence for the cell division process had a temporal resolution of 12 fpm.

For live-imaging of *hb*-MS2 signal, the pixel size remained at 132 nm, with *z*-step sizes adjusted to either 0.5 or 0.6 μm. These image sequences were acquired at a resolution of 512 × 512 pixels, with a temporal resolution set to 2 fpm. The excitation wavelengths used were 488 nm for MCP-GFP and 561 nm for Histone-RFP.

### Fixed imaging sample preparation and data acquisition

Data of 82 OreR embryos were from previous studies^35,42^. All other embryos were collected, fixed, and labeled according to a previously published protocol^35,42^. Briefly, *his2av-mrfp1* embryos collected at 25 °C were fixed in 8% (v/v) paraformaldehyde solution for 30 min, hand devitellinized, and stored in 1× PBS with 0.1% (w/v) BSA and 0.1% (v/v) Triton X-100 at 4 °C. OreR and 1×*bcd* embryos collected at 25 °C were fixed in 4% (v/v) paraformaldehyde solution for 15 min, vortexed in 100% methanol for devitellinization, and stored in 100% methanol at −20 °C. For OreR and 1×*bcd* embryos, *hb* or *Kr* mRNAs were labeled using smFISH, while Hb and Bcd proteins were labeled using immunofluorescence following smFISH (Bcd primary antibody: Santa Cruz Biotechnology, SC-66818; Hb primary antibody: Asian Distribution Center for Segmentation Antibodies, 576). The embryos’ nuclear DNA was stained subsequently with Hoechst 33342. *his2av-mrfp1* embryos were only stained for nuclear DNA. Following staining, embryos were washed (4 × 10 min) in PBTx (1× PBS, 0.1% (v/v) Triton X-100) and mounted in Aqua-Poly/Mount (Polysciences, 18606) for imaging.

Fixed embryos were imaged using a Zeiss LSM 880 confocal microscope equipped with a GaAsP detector and a 63×/1.4 NA oil immersion objective. Imaging of fixed *his2av-mrfp1* embryos used the same parameter setting as in live imaging. Multiple adjacent image stacks, with a typical size of ∼1700 × 1400 × 10 pixels each, were acquired to cover the cortex layer of each embryo. Imaging of fixed 1×*bcd* and OreR embryos was performed at 16-bit and 1 AU pinhole aperture with a pixel size of 66 nm and a *z*-step size of 0.32 μm. Multiple adjacent image stacks, with a typical size of ∼3200 × 3000 × 21 pixels each, were acquired to cover the cortex layer of each embryo.

### Image preprocessing and nuclear segmentation

Image processing followed a previously developed pipeline^35,42^ with updated algorithms. Briefly, raw images were converted to TIFF format and flat-field corrected. For both live and fixed images of Histone-RFP signals, two-dimensional (2D) nuclear segmentation was conducted on maximum intensity projection of the *z*-stack using a combination of local threshold and watershed. For fixed images of OreR and 1×*bcd* embryos, three-dimensional (3D) segmentation of nuclei was performed on Hoechst image stacks using the Cellpose algorithm^62^. Segmentation results were manually refined using a custom MATLAB graphical user interface (GUI). For each fixed embryo, the nuclear cycle was determined based on the number of recognized nuclei, using a criterium established from a study of 167 OreR embryos (Supplementary Fig. 2c). The embryo boundary was identified by thresholding the Hoechst image and was manually refined using custom MATLAB GUI. This boundary was used to determine the AP position of each nucleus.

### mRNA quantification in live images

In each frame of an image sequence, nascent mRNA foci candidates were identified as 3D local maxima in the nuclear region. Following a double-check using a custom MATLAB GUI, each focus was assigned to its closest nucleus. The local intensity profile of each focus was fitted to a 2D Gaussian function with a uniform background to extract the peak height (*I*_peak_) and radius (*σ*_0_), from which the fluorescence intensity of the focus was calculated as *I* = 2π*I*_peak_*σ*_0_^2^. By averaging foci intensities over all nuclei for each frame, the nuclear expression dynamics was extracted and normalized against its maximum value throughout the sequence. Results from multiple image sequences were further averaged and normalized.

### mRNA and protein quantification in fixed images

mRNA and protein quantification in fixed images followed established protocols^35,42^. Briefly, mRNA spot identification and intensity extraction were identical to those in live images. By comparing the joint distribution of peak height (*I*_peak_) and radius (*σ*_0_) between spots from the high-expression region (*hb*: anterior; *Kr*: medial) and those from a low-expression region (posterior), we determined a 2D threshold to distinguish real mRNA spots from background noise. The typical intensity, *I*_0_, of a single mRNA molecule was extracted from the primary peak of the spot intensity distribution based on a multi-Gaussian fitting. A threshold (3*I*_0_ for *hb* and *I*_0_ for *Kr*) was applied to identify active transcription sites within the recognized mRNA spots. The intensity of each transcription site was divided by *I*_0_ to determine the equivalent number of nascent transcripts at that site.

For Bcd and Hb protein signals, the average immunofluorescence intensity of each nucleus was estimated from the central *z*-slice of the nucleus. A background fluorescence level was estimated from the posterior part of the embryo and subtracted from the results. The typical intensity of a single protein molecule was extracted from cytoplasmic protein spots, similar to mRNA spot quantification. This value converted the background-subtracted nuclear fluorescence into absolute protein concentration.

### Determining the developmental time for live images

Each image sequence of a developing embryo covered multiple nuclear cleavage cycles, whose timing was determined based on nuclear division events. Specifically, each nuclear division event typically occurred in ∼1-2 min (nc10-11, nc11-12, and nc12-13: ∼1 min, nc13-14: ∼2 min). The last frame of each division event was defined as the start time of a cycle (Supplementary Fig. 1a). Since nuclear division event can be easily identified by eye, these mitotic frames were excluded from model training and prediction.

To align data from different embryos and compensate for the fluctuation in developmental tempo, we rescaled the nuclear cycle durations (excluding the mitotic frames) of each embryo to standard values. Specifically, in most embryos, nc11 interphase lasted ∼7 min, nc12 interphase lasted ∼8-10 min, and nc13 interphase lasted ∼12-15 min. We thus determined the standard durations of nc11-13 interphases to be 7, 9, and 13 min, respectively. In contrast, for the extended nc14 (lasting >1 hour), we focused on its early interphase (∼30 min), during which the existing time-prediction methods based on membrane invagination or nuclear length show limited accuracy^20,23^. We noticed that, during this period, nuclear size experienced a rapid increase followed by a gradual decline, stabilizing after ∼26 min (Supplementary Fig. 1d). Therefore, we rescaled the duration of early nc14 interphase based on this trend. To minimize the impact of imaging-induced phototoxicity, which could disrupt developmental timing, we excluded embryos with nuclear aberrations or significantly prolonged early nc14 interphase (≥30 min) from analysis.

Beyond embryo-to-embryo differences in cycle duration, alignment of different image sequences is further limited by the 1-minute resolution of time-lapse imaging, which can result in sub-minute time offsets between sequences. This offset is particularly critical in nc11, as its short duration means that even small timing differences can significantly impact nuclear morphology. To improve the accuracy of nc11 dataset labeling, we tried to extract additional temporal information from the first frame of nc11 in each image sequence. During this 1-minute interval, dividing nuclei undergo significant elongation, with the axial ratio changing systematically over time. By comparing the median nuclear axial ratio (*ε*%; major vs. minor axes) measured for each embryo, we classified all embryos into two groups with *ε*% around 1.36 and 1.67, respectively (Supplementary Fig. 1b). Using higher temporal resolution imaging (12 fpm) of nuclear division, we estimated the time offset between these two groups to be ∼25 s (∼0.4 min, Supplementary Fig. 1c). Consequently, nc11 time labels were adjusted to 0.6-6.6 min and 1-7 min for the two groups of embryos, accordingly.

### Generating datasets from live images for histone-based learning

For each nuclear cycle, live-imaging embryos were divided into two groups for training and testing, respectively (nc11: *n*_training_ = 13, *n*_test_ = 5; nc12: *n*_training_ = 12, *n*_test_ = 6; nc13: *n*_training_ = 18, *n*_test_ = 4; nc14: *n*_training_ = 5, *n*_test_ = 3). Within either group, the focal plane of each nucleus in every embryo was identified by comparing the nuclear histone signal in different *z*-slices. Histone images around each nucleus were cropped and normalized at the nuclear focal plane to generate three datasets of different spatial scales for training or testing, depending on the embryo group. For each nuclear cycle, the three spatial scales were adjusted to cover ∼1, 6, and 20 nuclei, i.e., nc11: 81 × 81, 301 × 301, and 601 × 601 pixels; nc12: 75 × 75, 251 × 251, and 401 × 401 pixels; nc13: 71 × 71, 201 × 201, and 401 × 401 pixels; nc14: 55 × 55, 151 × 151, and 251 × 251 pixels. All datasets were converted to 8-bit.

To test the impact of size reduction on time prediction, the image crop size of each dataset was increased (nc11: 90 × 90, 334 × 334, and 668 × 668 pixels; nc12: 83 × 83, 279 × 279, and 446 × 446 pixels; nc13: 79 × 79, 223 × 223, and 446 × 446 pixels) to mimic a 10% reduction of nuclear size.

### Generating datasets from fixed *his2av-mrfp1* images for relayed learning

160 fixed-imaging embryos covering every minute in nc11−13 (∼2 embryos/min) were used to perform relayed learning for DNA-based models. Firstly, histone images from each embryo were cropped and normalized around individual nuclei to generate nine datasets (corresponding to three spatial scales across three regions of the embryo) for histone-based time inference. Specifically, cropped images from the anterior (0.2 EL), medial (0.5 EL), and posterior (0.8 EL) regions of the fixed embryo (region sizes: 801 × 801 pixels for nc11, 1001 × 1001 pixels for nc12, and 1201 × 1201 pixels for nc13) were split into different datasets, with each region generating their own three-scale datasets. Due to fixation-induced embryo shrinkage, the image sizes of these datasets were rescaled to be: nc11: 66 × 66, 244 × 244, and 487 × 487 pixels; nc12: 62 × 62, 208 × 208, and 333 × 333 pixels; nc13: 59 × 59, 167 × 167, and 333 × 333 pixels (see **Image rescaling for time inference from fixed embryos**).

Secondly, the same set of embryos were divided into two groups for DNA-based training and testing, respectively (nc11: *n*_training_ = 16, *n*_test_ = 34; nc12: *n*_training_ = 20, *n*_test_ = 40; nc13: *n*_training_ = 22, *n*_test_ = 28). Within the training group, DNA images around each nucleus in every embryo were cropped and normalized to generate three datasets of different spatial scales, respectively, similar to the processing of live images. In contrast, DNA images from each embryo within the testing group were cropped and normalized around individual nuclei to generate nine datasets (corresponding to three spatial scales across three regions of the embryo), similar to the processing of fixed histone images. Due to fixation-induced embryo shrinkage, the image sizes of DNA datasets were adjusted to be: nc11: 70 × 70, 259 × 259, and 517 × 517 pixels; nc12: 66 × 66, 221 × 221, and 353 × 353 pixels; nc13: 63 × 63, 181 × 181, and 361 × 361 pixels. All datasets were converted to 8-bit.

### Generating datasets from fixed OreR and 1×*bcd* images for time prediction

Images and segmentation results of fixed OreR and 1×*bcd* embryos were resized to adjust the pixel dimensions to 132 × 132 nm^2^. The focal plane of each nucleus was identified based on the DNA signal. DNA images from each embryo were cropped and normalized around individual nuclei to generate nine datasets (corresponding to three spatial scales across three regions of the embryo) for DNA-based time prediction, similar to the processing of fixed histone images. Due to the difference in embryo size between strains, the image sizes of these datasets were rescaled. Specifically, the image sizes of OreR datasets were adjusted to be: nc11: 80 × 80, 296 × 296, and 590 × 590 pixels; nc12: 74 × 74, 248 × 248, and 397 × 397 pixels; nc13: 71 × 71, 203 × 203, and 406 × 406 pixels. Similarly, those of 1×*bcd* datasets were adjusted to be: nc11: 72 × 72, 267 × 267, and 533 × 533 pixels; nc12: 67 × 67, 226 × 226, and 360 × 360 pixels; nc13: 64 × 64, 185 × 185; and 368 × 368 pixels. All datasets were converted to 8-bit.

### Measuring the trend of nuclear diameter change over time

For every live or fixed images of *his2av-mrfp1* embryos, the diameter of each nucleus was measured, from which the median value was extracted. Results from different images were plotted against the measured or inferred developmental time and averaged with a bin size of 1 min (Supplementary Fig. 1d).

### Image rescaling for time inference from fixed embryos

To predict time from fixed *his2av-mrfp1* embryos using live-imaging-based models, the crop sizes for generating time inference datasets were rescaled to correct fixation-induced embryo shrinkage. The rescaling magnitude, *r*, for each nuclear cycle was determined through a recursion. Briefly, an initial guess of *r* = 1.18 was set based on the size comparison of fixed and live embryos. The crop sizes for three spatial scales were adjusted accordingly to generate tentative datasets for each embryo. Applying the trained histone model to these datasets predicted the developmental time of each embryo, which enabled reconstructing a trend of nuclear diameter change over time. Least-squares fitting of this reconstructed trend to live imaging result provided a new estimation of *r* for the next round of recursion. The convergence values of *r* (nc11: 1.23; nc12: 1.20; nc13: 1.20) were used to finalize the crop sizes for fixed *his2av-mrfp1* embryos.

Applying DNA-based models established from fixed *his2av-mrfp1* embryos on fixed OreR and 1×*bcd* embryos also requires rescaling the image crop sizes to account for embryo size difference between strains. Here, the rescaling magnitude, *r*, was estimated from the total area of the embryo. Specifically, the mean area of fixed *his2av-mrfp1* embryos was used as a standard (nc11: 5.7 × 10^4^ μm^2^, nc12−13: 5.8 × 10^4^ μm^2^). The mean area of fixed OreR and 1×*bcd* embryos (OreR: 7.4× 10^4^ μm^2^, 1×*bcd*: 6.1 × 10^4^ μm^2^) were measured as comparisons. For embryos whose areas fall within ±30% of the mean area, *r* was determined to be 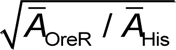 or 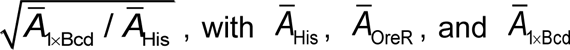 representing the mean areas of *his2av-mrfp1*, OreR, and 1×*bcd* embryos, respectively. For embryos whose areas fall between ±30% and ±50% of the mean area, *r* was determined based on the nearest ±30% boundary, i.e., 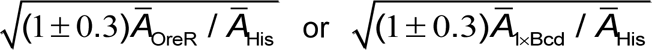. Embryos outside the ±50% mean area range were excluded as outliers.

### Calibrating image contrast and saturation for DNA images

Similar to fixed histone images, DNA signals from fixed embryos were also affected by embryo-to-embryo fluctuation in labeling and imaging. Impacting the background fluorescence level, this extra noise may affect the accuracy of our DNA-based models. To identify and solve this issue in each cropped DNA image, we first fitted the intensity histogram of the image to a multi-Gaussian distribution to extract the peak intensity (*I*_bk_) and width (*σ*_bk_) of background fluorescence. Measuring the median values of these quantities for each embryo revealed their upper limits, i.e., 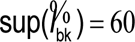 and 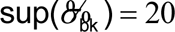 (Supplementary Figs. 3a-b). We thus increased the noise levels of each cropped image, i.e., lowered the image contrast, by adding a Gaussian white noise with *μ* = (60 − *I*_bk_)^+^and 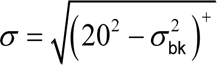.

Besides being affected by the background level, our DNA-based models were also sensitive to image saturation, which varied depending on the researcher’s imaging setting. Properly compensating for this effect is essential for integrating data from multiple experiments. To create a general framework for this task, we first quantified the saturation level (*r*_sat_) of each background-calibrated DNA image crop as the percentage of pixels at maximum intensity (255 for 8-bit images). By comparing the average *r*_sat_ of different training embryos, we classified all embryos into low-saturation and high-saturation groups with a threshold of 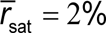 (Supplementary Fig. 3c). For each low-saturation image crop, we increased its saturation level to 2% via linear stretching of pixel intensities.

### CNN model construction, training, and testing

For each nuclear cycle, three independent CNN models for different image scales were constructed based on a modified VGGNet architecture^30^, with linear regression output (instead of softmax) for continuous-time inference. Besides the output form, the model was customized in the following ways to fit the small sample size of our study (thousands of images per dataset): (1) The number of convolutional layers was lowered from 8−16 to 3−6 to reduce overfitting. (2) Each convolutional layer was followed by a max-pooling layer to help reduce the parameter number. (3) The size of the convolutional kernel was increased from 3 × 3 to 5 × 5 to enhance feature mining with fewer layers. (4) The number of fully connected layers was increased from three to six to improve feature cross and classification, while the channel number for each layer was reduced to limit the parameter number. (5) In addition to dropout in the first two fully connected layers (dropout ratio: 50%), we added a dropout layer between the second and third fully connected layers (dropout ratio: ∼50%) in some CNN models to reduce overfitting. (6) A batch normalization layer was inserted before the flatten layer to speed up model training and reduce overfitting. (7) Based on the characteristics of fly embryo images, we chose to perform random rotation (0−30°), horizontal flip, and a slight shear transformation (0−0.1) in data augmentation. Specifically, the shear transformation was applied to account for slight nuclear deformation that may happen during the experiment. The detailed information of all models is listed in Supplementary Table 1.

For model training, thousands of cropped images from the corresponding training dataset were resized and converted to normalized float format for input. Mean Square Error (MSE) and Adaptive Moment Estimation (ADAM) were selected as the loss function and the optimization function, respectively, with a learning rate of 0.001. With *k*-fold cross-validation (*k* = 5 or 10) and Mean Absolute Error (MAE)-based evaluation metric, each model was trained for ∼400 epochs to find the best result. Following the above protocol, we adjusted the image batch size, the depth and width of convolutional layers, and the number and rate of dropout layers for each model to further optimize model parameters.

To test the performance of a model on an embryo (or its anterior, medial, or posterior region), cropped images from the corresponding test dataset were resized and converted to normalized float format for input. Typically, the raw image stack of an embryo can generate hundreds of small- and median-scale images and >40 large-scale images, each of which can provide a time prediction. To balance prediction speed and accuracy, we randomly selected 40 images of the corresponding scale and recorded the median value of individual images’ prediction results as the model’s time inference for the embryo (or its anterior, medial, or posterior region).

Model training and testing were performed on a workstation equipped with an NVIDIA GeForce RTX 3090 GPU. Training each model took 6−24 h.

### Integration of time prediction results

For each embryo in the test group, time prediction results from three independent CNN models for different image scales were compared and integrated. Specifically, for a live-imaging embryo, whose images only covered a small region of the embryo, the median value of three independent predictions from this imaged region was output as the final inference result. Conversely, for a fully imaged fixed embryo, the anterior, medial, and posterior regions each provided three independent predictions. Each region’s median prediction was extracted and compared with each other. An average over the three regions was output as the inference result for the embryo. In addition, for DNA-based inference, the standard deviation of all nine predictions (*σ*_infer_) served as a confidence criterion (Supplementary Fig. 3d). Embryos with *σ*_infer_ > 1.5 min were discarded for further analysis.

### Measuring the spatial expression patterns of Bcd, Hb, and *Kr*

For each OreR and 1×*bcd* embryo labeled with *Kr* mRNA and Bcd/Hb proteins, we plotted the nascent mRNA signal (*r*, in units of the number of molecules) and absolute protein concentrations against nuclear position (*x*) for all nuclei, respectively. For each data species, we binned individual data points by *x*. Within position ranges 0.10−0.75 EL (for Bcd), 0.15−0.85 EL (for Hb), 0.20−0.85 EL (for *Kr* in OreR), and 0.10−0.80 EL (for *Kr* in 1×*bcd*), we used a least-squares algorithm (the “fit” function in MATLAB) to fit the binned data from each species to the following functions:

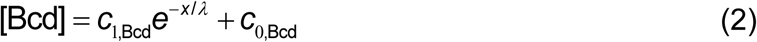

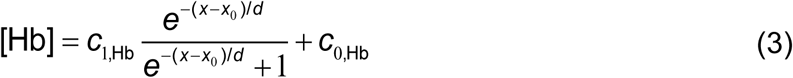

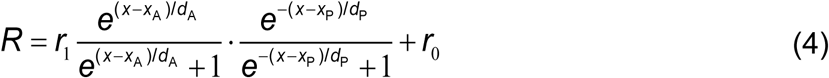

where *c*_1,·_, *c*_0,·_, *r*_1_, and *r*_0_ denote the peak and background levels of the corresponding species, respectively; *λ*, *d*, and *d*_A_ indicate the decay lengths of Bcd, Hb, and *Kr* patterns, respectively; *x*_0_ represents the boundary position of Hb expression pattern; *x*_A_ and *x*_P_ denote the anterior and posterior boundary positions of *Kr* expression domain, respectively. For *Kr*-boundary analysis (Supplementary Fig. 4d), low-expression embryos with <3 nascent *Kr* mRNAs per nucleus within the expression band (OreR: 0.4−0.6 EL, 1×*bcd*: 0.35−0.55 EL) were excluded.

### Thermodynamic modeling of unsteady-state gene regulation

To analyze the dynamic regulation of *Kr* by Bcd and Hb, we related the average nuclear expression of *Kr* to Bcd and Hb concentrations at each position within 0.20–0.70 EL for each embryo. By predicting the developmental time of each embryo, we binned the *Kr*-Bcd-Hb relationship at each spatial position along the time axis with a bin width of 1 min. To understand this spatiotemporal relationship, we constructed a time-dependent thermodynamic model, with *Kr* transcription determined by four possible states of cooperative Bcd and Hb bindings, i.e.,

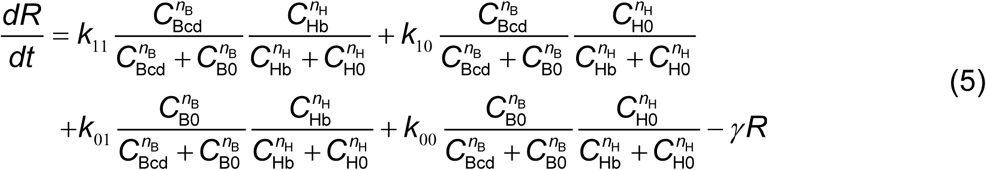

Here, the cooperativity of either Bcd or Hb is described by a Hill function of protein concentration, with *n*. and *C*._0_ representing the Hill coefficient and concentration threshold of the corresponding species, respectively. The mRNA production rate for each binding state is denoted by *k_ij_*, with *i*, *j* = 0 or 1 corresponding to the unbound or bound states of Bcd and Hb, respectively. Following production, mRNA is thought to degrade at a constant rate *γ*. By comparing the quantitative relationship between *Kr*, Bcd, and Hb to Equation (5), we determined that only *k*_10_ is non-zero, leading to Equation (1).

Based on the observed *Kr* expression dynamics, we assumed that the gene is only activatable within a specific period in each nuclear cycle. For nc11−12, we used two parameters, *t*_on_ and *t*_off_, to represent the start and end moment of this period, respectively. For nc13, we incorporated two additional parameters, *t*_low,1_ and *t*_low,2_, to delineate the start and end moment of the observed low-expression phase within the activatable period, respectively. The transcription rate was considered to decrease by a factor *r* in this phase.

Following the above settings, we can numerically solve Equation (1) at each position and time point of a nuclear cycle based on the observed Bcd and Hb concentration profiles. Experimental data from different positions and time points were fitted altogether to theoretical results using a least-squares algorithm (the “lsqcurvefit” function in MATLAB) to extract kinetic parameters. The fitted time parameters (*t*_on_, *t*_off_, *t*_low,1_, *t*_low,2_) were used to extract five expression periods (*k* > 0) in nc11-13 (4.6-5.5 min in nc11; 4.8-6.6 min in nc12; 3.8-5 min, 5-8.5 min, and 8.5-11.5 min for nc13).

### PWM motif analysis

Sequences of the two *Kr* enhancers were provided in the reference^45^. Bcd and Hb binding sites on *Kr* enhancers were predicted using a PWM motif analysis following previous literature^57^. PWMs for Bcd and Hb were retrieved from the OnTheFly database^63^ (http://bhapp.c2b2.columbia.edu/OnTheFly/index.php). By computing the binding probabilities of Bcd and Hb at each location on the sequences, four strong Bcd sites and seven strong Hb sites were identified on *Kr* enhancers with the criterion: sites with a log-odds of above 5.0 (natural logarithm) (Supplementary Fig. 4e).

### Mathematical modeling of unsteady-state transcriptional kinetics

Stochastic modeling and inference of unsteady-state transcriptional kinetics are described in detail in Supplementary Note.

## Data availability

The raw image data reported in this paper are publicly accessible at a private server (http://gofile.me/4yuzx/XhyORolMN). Source data are provided with this paper.

## Code availability

Custom scripts for this paper were written in multiple programming languages and are available on GitHub (https://github.com/Xulab-biophysics/FISHIF-Time2024.git) and a private server (http://gofile.me/4yuzx/3MbMTb7H2). Specifically, scripts for image preprocessing, gene regulation analysis, and theoretical modeling were developed in MATLAB 2023a (MathWorks). CNN models for time inference were implemented using a Python-based deep learning API, Keras (https://keras.io/). A ready-to-used DNA-based time inference framework was encapsulated into a MATLAB APP for customer usage. PWM motif analysis was conducted using C++.

## Acknowledgments

We thank Hernan H. Garcia and Jun Ma for the generous gift of fly lines. We thank Asian Distribution Center for Segmentation Antibodies for providing the anti-Hb antibody. We thank Jingyao Wang for providing previously published imaging data. This work was supported by the National Key R&D Program of China (grant no. 2021YFA0910702), the National Natural Science Foundation of China (grant no. 12474194, 11774225, 41921006), and the Natural Science Foundation of Shanghai (grant no. 22ZR1434000). We gratefully acknowledge the imaging and computing resources provided by the Student Innovation Center at Shanghai Jiao Tong University, and we sincerely thank Liuyin Fan for the dedicated management and support of these resources.

## Author contributions

Conceptualization by H.B. and H.X.; Methodology by H.B., S.Z., and H.X.; Software by H.B., S.Z. and Z.Y.; Formal Analysis by H.B. and S.Z.; Investigation by H.B., S.Z., and H.X.; Writing – Original Draft by H.B., S.Z., and H.X.; Funding acquisition by H.X.; Resources by H.X.; and Supervision by H.X.

## Competing interests

The authors declare no competing interests.

## Notes

### Competing Interest Statement

The authors have declared no competing interest.

