## Supplementary Information for "Deep Learning-Based High-Resolution Time Inference for Deciphering Dynamic Gene Regulation from Fixed Embryos"

**Supplementary Note 1:** Mathematical modeling of unsteady-state transcription kinetics.

**Supplementary Fig. 1.** Training and testing the histone-based time predictor.

**Supplementary Fig. 2.** Adapting the histone-based time inference to fixed embryos.

**Supplementary Fig. 3.** Training and testing the DNA-based time predictor for fixed WT embryos.

**Supplementary Fig. 4.** Quantitative analysis of dynamic *Kr* regulation using fixed embryos.

**Supplementary Fig. 5.** Estimating kinetic parameters of the unsteady-state *hb* transcription model.

**Supplementary Table 1.** Configuration of CNN models.

**Supplementary References**

### Supplementary Note 1: Mathematical modeling of unsteady-state transcription kinetics.

#### 1.1. Model description

Previous studies have shown that the *hb* gene has two differentially expressed promoters, P1 and P2, each producing an alternative transcript of distinct length[1-4]. In early development, most transcripts are produced by the P2 promoter following a bursty stochastic process[4-7]. According to recent research, this bursty activity satisfies a three-state telegraph model, where the promoter randomly transitions between one inactive and two active states[2]. However, since the transition to one of the active states is rare, this model has often been simplified to a two-state version in prior studies[6,8].

This study modeled nascent transcription of a single hb gene locus as a time-dependent two-state process, considering gene shutdown and re-expression during each mitosis. Specifically, the gene is assumed to be silent at the beginning and end of each nuclear cycle and only activatable in the middle of the nuclear cycle. To account for variability in the timing of these periods between individual gene loci, we introduced a time-dependent variable, $P_{\text{active}}$, to describe the percentage of activatable gene loci at each time point. Since the mismatch between different loci's activatable periods is typically small compared to their overlap, $P_{\text{active}}(t)$ is sufficient to describe the statistical properties of individual loci's silent and activatable periods. Typically, $P_{\text{active}}$ varies in three phases within each nuclear cycle: increasing from zero to its maximum at the beginning, staying stable through the middle, and decreasing back to zero at the end.

During an activatable period, the gene randomly switches between two transcription states: an "OFF" state (denoted as state 0), where the gene is inactive, and an "ON" state (denoted as state 1), where the gene actively initiates new transcripts. State transitions and mRNA initiation are assumed to be Poisson processes with time-dependent rates $k_{ij}$ and $k_{\mathrm{INI},i}$ (where $i, j$ = 0, 1)[2,5,6,8]. In this study, $k_{01}$, $k_{10}$, and $k_{\mathrm{INI},1}$ are renamed $k_{\mathrm{ON}}$, $k_{\mathrm{OFF}}$, and $k_{\mathrm{INI}}$, respectively, while $k_{\mathrm{INI},0}$ is set to zero. Following initiation, each nascent mRNA molecule elongates to its final length $L$ with a constant speed $V_{\mathrm{EL}}$, and then stays on the gene for an extra termination period, $T_{\mathrm{R}}$, before release[2,7,8]. During the silent period, the gene remains in the "OFF" state.

At a given observation time $t_{\mathrm{ob}}$, the state of the system is determined by the gene state $n$ ($n$ = 0, 1) and the total signal of nascent mRNA $m$ ($m \geq 0$). Following previous *hb* kinetics models[2,6], $m$ is computed as the sum of signals from all transcripts initiated within a fixed period $T_{\mathrm{RES}} = L/V_{\mathrm{EL}} + T_{\mathrm{R}}$ before $t_{\mathrm{ob}}$, i.e., $m = \sum_{-T_{\mathrm{RES}} \leq \tau_i \leq 0} g(\tau_i)$. Here, $\tau_i = t_i - t_{\mathrm{ob}}$ is the initiation time relative to $t_{\mathrm{ob}}$, and the signal from a single nascent transcript is described by a contribution function $g(\tau)$, whose shape depends on the target positions of the probe set and the magnitude of $T_{\mathrm{R}}$[2,7,9]. In this paper, we used $L$ = 3621 bp and $T_{\mathrm{R}}$ = 0, based on previous studies[2,8], while leaving $V_{\mathrm{EL}}$ as a variable to be determined.

### 1.2. Master equation

We may first model the behavior of a single gene locus during the activatable period, with the start and end times denoted as $t_{\text{start}}$ and $t_{\text{end}}$, respectively. The master equation for computing the probability distribution of ($n$, $m$) at $t_{\text{ob}}$ ($\tau = 0$) is written as

$$\frac{d\mathbf{P}(\mathscr{m})}{d\tau} = (\mathbf{K} - \mathbf{K}_{\text{INI}})\mathbf{P}(\mathscr{m}) + \mathbf{K}_{\text{INI}}\mathbf{P}(\mathscr{m} - g(\tau)) \tag{S1}$$

where $\mathscr{m}(\tau) = \sum_{-T_{\text{RES}} \le \tau_i \le \tau} g(\tau_i)$ is a pseudo-observable describing the accumulation of $m$ over the history from $t_{\text{ob}} - T_{\text{RES}}$ to $t_{\text{ob}} - \tau$, $\mathbf{P}(\mathscr{m}) = \begin{bmatrix} P(0,\mathscr{m}) \\ P(1,\mathscr{m}) \end{bmatrix}$ is a vectorized form of $P(n,\mathscr{m})$, $\mathbf{K} = \begin{bmatrix} -k_{\text{ON}} & k_{\text{OFF}} \\ k_{\text{ON}} & -k_{\text{OFF}} \end{bmatrix}$ and $\mathbf{K}_{\text{INI}} = \begin{bmatrix} 0 & 0 \\ 0 & k_{\text{INI}} \end{bmatrix}$ are matrices describing gene-state transition and transcription initiation, respectively[2,6,7]. Since $\mathscr{m} = 0$ for $\tau = -T_{\text{RES}}$ and $\mathscr{m} = m$ for $\tau = 0$, the distribution of ($n$, $m$) at $t_{\text{ob}}$ can be obtained by solving **Equation (S1)** with the initial condition $\mathbf{P}_{\tau=-T_{\text{RES}}}(\mathscr{m}) = \mathbf{q}\delta_{\mathscr{m},0}$, where $\mathbf{q}$ is the marginal distribution of the gene state at $\tau = -T_{\text{RES}}$, i.e., $\mathbf{q} = \begin{bmatrix} P_{n=0} \\ P_{n=1} \end{bmatrix}_{\tau=-T_{\text{RES}}}$. Formally, the solution of **Equation (S1)** is written as the propagation of the initial condition[2,6,7], i.e.,

$$\bar{\mathbf{P}}_{t_{\text{ob}}} = \mathbf{U}(0, -T_{\text{RES}})\bar{\mathbf{q}} \tag{S2}$$

where $\bar{\mathbf{P}}_{t_{\text{ob}}} \equiv \left[\mathbf{P}_{t_{\text{ob}}}(m)\right]^{\text{T}}$ and $\bar{\mathbf{q}} \equiv \left[\delta(\mathscr{m})\mathbf{q}\right]^{\text{T}}$ combine probabilities of different $m$ and $\mathscr{m}$ values into full vector forms, and $\mathbf{U}(0, -T_{\text{RES}}) = \mathcal{T}\left[\exp\left(\int_{-T_{\text{RES}}}^{0} \left(\bar{\mathbf{K}} + \bar{\mathbf{K}}_{\text{INI}}(\tau)\right) d\tau\right)\right]$ is the propagator from $\tau = -T_{\text{RES}}$ to $\tau = 0$. Here, $\mathcal{T}$ represents the time-ordering operator, while $\bar{\mathbf{K}} = \left[\mathbf{K}\delta(m - \mathscr{m})\right]$ and $\bar{\mathbf{K}}_{\text{INI}} = \left[\mathbf{K}_{\text{INI}}\left(\delta(m - \mathscr{m} - g(\tau)) - \delta(m - \mathscr{m})\right)\right]$ denote the full matrix version of **K** and $\mathbf{K}_{\mathrm{INI}}$ combining different $m$ and $\mathscr{m}$ values.

Notably, the above solution only applies when the entire propagation occurs within the activatable period, i.e., $[t_{\mathrm{ob}} - T_{\mathrm{RES}}, t_{\mathrm{ob}}] \subset [t_{\mathrm{start}}, t_{\mathrm{end}}]$. If $t_{\mathrm{ob}} - T_{\mathrm{RES}} < t_{\mathrm{start}}$, the propagation should be modified as:

$$\bar{\mathbf{P}}_{t_{\mathrm{ob}}} = \mathbf{U}(0, t_{\mathrm{start}} - t_{\mathrm{ob}})\bar{\mathbf{q}}_{\mathrm{start}} \tag{S3}$$

with $\mathbf{q}_{\mathrm{start}} = \begin{bmatrix} 1 \\ 0 \end{bmatrix}$ representing the marginal distribution of the gene state at $t_{\mathrm{start}}$. Similarly, if $t_{\mathrm{ob}} > t_{\mathrm{end}}$, the propagation should be modified as

$$\bar{\mathbf{P}}_{t_{\mathrm{ob}}} = \mathbf{U}(t_{\mathrm{end}} - t_{\mathrm{ob}}, T_{\mathrm{RES}})\bar{\mathbf{q}} \tag{S4}$$

where **q** denotes the marginal distribution of the gene state at $t_{\mathrm{ob}} - T_{\mathrm{RES}}$.

To describe the probability distribution across an ensemble of gene loci, we need to account for the effect of $P_{\mathrm{active}}$ in the above equations. Specifically, we can split $\mathbf{P}(\mathscr{m})$ into two parts, $\mathbf{P}_{\mathrm{active}}(\mathscr{m})$ and $\mathbf{P}_{\mathrm{inactive}}(\mathscr{m})$, to describe loci within and outside the activatable period, respectively. Their time evolutions are interrelated as follows:

$$\begin{cases} \dfrac{d\mathbf{P}_{\mathrm{active}}(\mathscr{m})}{d\tau} = (\mathbf{K} - \mathbf{K}_{\mathrm{INI}})\mathbf{P}_{\mathrm{active}}(\mathscr{m}) + \mathbf{K}_{\mathrm{INI}}\mathbf{P}_{\mathrm{active}}(\mathscr{m} - g(\tau)) - k_{\mathrm{i}}\mathbf{P}_{\mathrm{active}} + k_{\mathrm{a}}\mathbf{P}_{\mathrm{inactive}} \\ \dfrac{d\mathbf{P}_{\mathrm{inactive}}(\mathscr{m})}{d\tau} = k_{\mathrm{i}}\mathbf{P}_{\mathrm{active}} - k_{\mathrm{a}}\mathbf{P}_{\mathrm{inactive}} \end{cases} \tag{S5}$$

where $k_{\mathrm{a}} = \left(\dfrac{\dot{P}_{\mathrm{active}}(\tau)}{P_{\mathrm{active}}(\tau)}\right)^{+}$ and $k_{\mathrm{i}} = -\left(\dfrac{\dot{P}_{\mathrm{active}}(\tau)}{P_{\mathrm{active}}(\tau)}\right)^{-}$ denote the relative increasing and decreasing rates of $P_{\mathrm{active}}$, respectively. Similarly, the initial condition **q** is divided into

$\mathbf{q}_{\text{active}} = \begin{bmatrix} P_{\text{active}} - P_{n=1} \\ P_{n=1} \end{bmatrix}_{\tau=-T_{\text{RES}}}$ and $\mathbf{q}_{\text{inactive}} = \begin{bmatrix} 1 - P_{\text{active}} \\ 0 \end{bmatrix}_{\tau=-T_{\text{RES}}}$ to describe the marginal distributions of gene states for loci within and out of the activatable period, respectively. The overall ensemble probability distribution of ($n$, $m$) is then solved for three different cases, respectively, i.e.:

(1) If $P_{\text{active}}$ remains constant during the propagation period ([$t_{\text{ob}} - T_{\text{RES}}$, $t_{\text{ob}}$]),

$$\bar{\mathbf{P}}_{t_{\text{ob}}} = \mathbf{U}(0, -T_{\text{RES}})\bar{\mathbf{q}}_{\text{active}} + \bar{\mathbf{q}}_{\text{inactive}} \tag{S6}$$

(2) If $P_{\text{active}}$ increases during the propagation period,

$$\bar{\mathbf{P}}_{t_{\text{ob}}} = \mathbf{U}(0, -T_{\text{RES}})\bar{\mathbf{q}}_{\text{active}}(t_{\text{ob}} - T_{\text{RES}}) - \int_{-T_{\text{RES}}}^{0} \mathbf{U}(0, \tau)\dot{\bar{\mathbf{q}}}_{\text{inactive}}(t_{\text{ob}} + \tau)d\tau + \bar{\mathbf{q}}_{\text{inactive}}(t_{\text{ob}}) \tag{S7}$$

(3) If $P_{\text{active}}$ decreases during the propagation period,

$$\begin{aligned} \bar{\mathbf{P}}_{t_{\text{ob}}} = {} & \frac{P_{\text{active}}(t_{\text{ob}})}{P_{\text{active}}(t_{\text{ob}} - T_{\text{RES}})} \mathbf{U}(0, -T_{\text{RES}})\bar{\mathbf{q}}_{\text{active}}(t_{\text{ob}} - T_{\text{RES}}) \\ & + \int_{-T_{\text{RES}}}^{0} \frac{\dot{P}_{\text{active}}(t_{\text{ob}} + \tau)}{P_{\text{active}}(t_{\text{ob}} - T_{\text{RES}})} \mathbf{U}(\tau, -T_{\text{RES}})\bar{\mathbf{q}}_{\text{active}}(t_{\text{ob}} - T_{\text{RES}})d\tau + \bar{\mathbf{q}}_{\text{inactive}}(t_{\text{ob}} - T_{\text{RES}}) \end{aligned} \tag{S8}$$

#### 1.3. Numerically solving the master equation

We numerically implemented **Equations (S6)-(S8)** using the finite state projection (FSP) method[2,6,10,11]. Briefly, we discretized and truncated the range of $m$ to a finite set {0, $\Delta m$, $2\Delta m$, …, $m_{\max}$}, with $\Delta m \ll 1$ and $m_{\max}$ sufficiently large to capture the majority of the nascent mRNA distribution. Equation (S1) is then transformed into a finite-dimensional version:

$$\dot{\bar{\mathbf{P}}} = \left(\bar{\mathbf{K}} + \bar{\mathbf{K}}_{\text{INI}}(\tau)\right)\bar{\mathbf{P}} = \begin{bmatrix} \mathbf{K}-\mathbf{K}_{\text{INI}} & 0 & 0 & 0 \\ 0 & \mathbf{K}-\mathbf{K}_{\text{INI}} & 0 & \cdots \\ 0 & 0 & \mathbf{K}-\mathbf{K}_{\text{INI}} & \cdots \\ \vdots & \vdots & \vdots & \ddots \\ \mathbf{K}_{\text{INI}} & 0 & 0 & \ddots \\ 0 & \mathbf{K}_{\text{INI}} & 0 & \ddots \\ 0 & 0 & \mathbf{K}_{\text{INI}} & \ddots \\ \vdots & \vdots & \vdots & \ddots \end{bmatrix} \begin{bmatrix} \mathbf{P}(0) \\ \mathbf{P}(\Delta m) \\ \mathbf{P}(2\Delta m) \\ \vdots \\ \mathbf{P}(g(\tau)) \\ \mathbf{P}(g(\tau)+\Delta m) \\ \mathbf{P}(g(\tau)+2\Delta m) \\ \vdots \end{bmatrix} \tag{S9}$$

where $\bar{\mathbf{K}} = \begin{bmatrix} \mathbf{K} & 0 & 0 & \cdots \\ 0 & \mathbf{K} & 0 & \cdots \\ 0 & 0 & \mathbf{K} & \cdots \\ \vdots & \vdots & \vdots & \ddots \end{bmatrix}$ and $\bar{\mathbf{K}}_{\text{INI}}(\tau) = \begin{bmatrix} -\mathbf{K}_{\text{INI}} & 0 & 0 & \cdots \\ 0 & -\mathbf{K}_{\text{INI}} & 0 & \cdots \\ \vdots & 0 & -\mathbf{K}_{\text{INI}} & \cdots \\ \mathbf{K}_{\text{INI}} & \vdots & 0 & \cdots \\ 0 & \mathbf{K}_{\text{INI}} & \vdots & \ddots \\ 0 & 0 & \mathbf{K}_{\text{INI}} & \ddots \\ \vdots & \vdots & \vdots & \ddots \end{bmatrix}$.

By further discretizing the range of propagation time into a finite series {$-T_{\text{RES}}$, $\Delta\tau - T_{\text{RES}}$,…, $-2\Delta\tau$, $-\Delta\tau$, 0} with $\Delta\tau \ll T_{\text{RES}}$, we rewrote Equation (S6) as:

$$\begin{aligned} \bar{\mathbf{P}}(t_{\text{ob}}) &= \mathbf{U}(0,-\Delta\tau)\cdots\mathbf{U}(\Delta\tau - T_{\text{RES}}, -T_{\text{RES}})\bar{\mathbf{q}}_{\text{active}}(t_{\text{ob}} - T_{\text{RES}}) + \bar{\mathbf{q}}_{\text{inactive}}(t_{\text{ob}} - T_{\text{RES}}) \\ &= \left(\mathbf{I} + \bar{\mathbf{K}}\Delta\tau + \bar{\mathbf{K}}_{\text{INI}}(-\Delta\tau)\Delta\tau\right)\cdots\left(\mathbf{I} + \bar{\mathbf{K}}\Delta\tau + \bar{\mathbf{K}}_{\text{INI}}(-T_{\text{RES}})\Delta\tau\right)\bar{\mathbf{q}}_{\text{active}}(t_{\text{ob}} - T_{\text{RES}}) \\ &\quad + \bar{\mathbf{q}}_{\text{inactive}}(t_{\text{ob}} - T_{\text{RES}}) \end{aligned} \tag{S10}$$

where $\mathbf{U}(\tau+\Delta\tau,\tau) = \left(\mathbf{I} + \bar{\mathbf{K}}\Delta\tau + \bar{\mathbf{K}}_{\text{INI}}(\tau)\Delta\tau\right)$ is the infinitesimal propagator from time $\tau$ to $\tau + \Delta\tau$, and **I** is the unit matrix.

Equations (S7) and (S8) are handled similarly, except that the variation of $P_{\text{active}}$ needs to be considered in the expression. Specifically, Equation (S7) is rewritten as:

$$\bar{\mathbf{P}}(t_{ob}) = \mathbf{U}(0,-\Delta\tau)\left(\cdots\left(\mathbf{U}(\Delta\tau - T_{RES}, -T_{RES})\bar{\mathbf{q}}_{active}(t_{ob} - T_{RES}) - \Delta\bar{\mathbf{q}}_{inactive}(t_{ob} - T_{RES})\right)\cdots\right) \\ -\Delta\bar{\mathbf{q}}_{inactive}(t_{ob} - \Delta\tau) + \bar{\mathbf{q}}_{inactive}(t_{ob}) \tag{S11}$$

Where $\Delta\bar{\mathbf{q}}_{inactive}(t) = \bar{\mathbf{q}}_{inactive}(t + \Delta\tau) - \bar{\mathbf{q}}_{inactive}(t) = -(P_{active}(t + \Delta\tau) - P_{active}(t))\bar{\mathbf{q}}_{start}$.

**Equation (S8)** is rewritten as:

$$\begin{aligned} \bar{\mathbf{P}}(t_{ob}) = & \frac{P_{active}(t_{ob})}{P_{active}(t_{ob} - T_{RES})}\mathbf{U}(0,-\Delta\tau)\cdots\mathbf{U}(\Delta\tau - T_{RES}, -T_{RES})\bar{\mathbf{q}}_{active}(t_{ob} - T_{RES}) \\ & + \frac{\Delta P_{active}(t_{ob} - \Delta\tau)}{P_{active}(t_{ob} - T_{RES})}\mathbf{U}(-\Delta\tau, -2\Delta\tau)\cdots\mathbf{U}(\Delta\tau - T_{RES}, -T_{RES})\bar{\mathbf{q}}_{active}(t_{ob} - T_{RES}) \\ & + \cdots \\ & + \frac{\Delta P_{active}(t_{ob} - T_{RES})}{P_{active}(t_{ob} - T_{RES})}\mathbf{U}(\Delta\tau - T_{RES}, -T_{RES})\bar{\mathbf{q}}_{active}(t_{ob} - T_{RES}) \\ & + \bar{\mathbf{q}}_{inactive}(t_{ob}) \end{aligned} \tag{S12}$$

where $\Delta P_{active}(t) = P_{active}(t + \Delta\tau) - P_{active}(t)$. In practice, all terms but the last one on the right-hand side of the equation can be computed together through a single propagation.

In this paper, we used $\Delta m$ = 1 and $\Delta\tau$ = 0.1 s to balance computational accuracy and speed. It is important to note that $k_{ON}$, $k_{OFF}$, $k_{INI}$, and $P_{active}$ are all time dependent and may vary during the propagation time window ([$t_{ob} - T_{RES}$, $t_{ob}$]). If two observations are performed at closely spaced time points, i.e., $\Delta t_{ob} < T_{RES}$, their propagation time windows will partially overlap, during which the corresponding kinetic parameters and gene states for the two propagations will be identical. I.e., the latter propagation inherits kinetic parameters and gene states from the former one.

#### 1.4. Modeling the DNA replication effect

Due to gene replication during each nuclear cycle, some of the observed *hb* foci may correspond to pairs of closely located sister loci that are indistinguishable under the microscope[2,5,8]. To account for this effect, we followed previous literature to assume that the two sister gene copies are expressed independently[2,5,8]. Supposing a certain percentage ($\alpha$) of the observed *hb* foci are from indistinguishable sister loci pairs, we wrote the distribution of the observed nascent mRNAs ($m_{ob}$) at individual foci as

$$P(m_{ob}) = (1-\alpha)P(m_{single}) + \alpha P(m_{single}) * P(m_{single}) \quad \text{(S13)}$$

where $P(m_{single})$ denotes the nascent mRNA distribution of a single gene copy computed from the model, and "$*$" indicates convolution.

#### 1.5. Inferring the transcription kinetics

We inferred the unsteady-state *hb* kinetics from time-resolved experimental data using a modified maximum likelihood estimation (MLE) method[6,11]. Briefly, based on the predicted embryonic times, we grouped all embryos within each nuclear cycle into multiple 1-minute time windows. For each time window, which typically contains ≥3 embryos, we computed $t_{ob}$ as the average developmental time and pooled the single-locus nascent mRNA data from all embryos into multiple nuclear position bins. To ensure sufficient data points in each bin, we used overlapping binning with a bin size of 0.1 EL.

Since $T_{\mathrm{RES}}$ for *hb* is believed to be greater than 1 minute[12,13], the nascent mRNA distributions in neighboring time windows are correlated. Therefore, for each nuclear position bin, we fitted data from all time windows together. Specifically, for a given time-dependent parameter set $\tilde{\mathbf{K}} = \{k_{\mathrm{ON}}(t_{\mathrm{ob}}), k_{\mathrm{OFF}}(t_{\mathrm{ob}}), k_{\mathrm{INI}}(t_{\mathrm{ob}}), P_{\mathrm{active}}(t_{\mathrm{ob}}), \alpha(t_{\mathrm{ob}}), V_{\mathrm{EL}}\}$, the likelihood of observing a time-resolved data set is

$$L(M \mid \tilde{\mathbf{K}}) = \prod_{i,j} P(m_i(t_{\mathrm{ob},j}) \mid \tilde{\mathbf{K}}) \tag{S14}$$

where $M = \{m_i(t_{\mathrm{ob},j})\}$ is the single-locus data set from all time windows (each with a different $t_{\mathrm{ob}}$) and $P(m_i(t_{\mathrm{ob},j}) \mid \tilde{\mathbf{K}})$ is the probability of observing $m_i$ nascent mRNAs at the $j$-th $t_{\mathrm{ob}}$, given $\tilde{\mathbf{K}}$. Note that all parameters but $V_{\mathrm{EL}}$ in $\tilde{\mathbf{K}}$ are time dependent.

To efficiently compute the nascent mRNA distribution for each $t_{\mathrm{ob}}$ in **Equation (S14)**, we started from the first time point, where the initial condition for propagation is set to be $\mathbf{q}_{\mathrm{active}} = \begin{bmatrix} 0 \\ 0 \end{bmatrix}$ and $\mathbf{q}_{\mathrm{inactive}} = \begin{bmatrix} 1 \\ 0 \end{bmatrix}$ at $t = t_{\mathrm{ob},1} - T_{\mathrm{RES}}$. The initial conditions for subsequent time points $t_{\mathrm{ob},j}$ are determined iteratively, using the marginal distributions of the gene state at $t = t_{\mathrm{ob},j} - T_{\mathrm{RES}}$, which can be obtained from the propagation for $t_{\mathrm{ob},j-1}$. During each propagation, since $\Delta t_{\mathrm{ob}}$ is much larger than the propagation time step, we treated all time-dependent parameters as step functions being stable within $\Delta t_{\mathrm{ob}}$.

We performed a parameter search to maximize the likelihood over a broad range of values ($\alpha$ from 0 to 1, $P_{\mathrm{active}}$ from 0 to 1, $k_{\mathrm{ON}}$ and $k_{\mathrm{OFF}}$ from 0 to 10 min$^{-1}$, $k_{\mathrm{INI}}$ from 0 to 100 min$^{-1}$, $V_{\mathrm{EL}}$ from 25 to 55 bp/s) using a combination of simplex and simulated annealing methods. Based on the fitting results for the anterior nuclear position bin (0.2-0.4 EL), where *hb* is highly expressed, we first determined that $V_{EL}$ = 40 bp/s (**Supplementary Fig. 5a**). With this value fixed, we then analyzed the spatiotemporal profiles of other parameters. Specifically, we found that $P_{active}$ showed similar temporal profiles across different nuclear position bins (**Fig. 5c**). Thus, we established a universal $P_{active}$ profile for the entire embryo by averaging the profiles extracted from all nuclear positions. Moreover, we observed that $k_{ON}$ varied significantly with nuclear position and time, while $k_{OFF}$ and $k_{INI}$ remained stable (**Supplementary Figs. 5b-c**). Therefore, we fixed $k_{OFF}$ and $k_{INI}$ at their mean values before conducting a detailed re-scan of $k_{ON}$. To increase the accuracy of simplex and simulated annealing methods in the above steps, we repeated each search 12 times and selected the result with the highest likelihood.

Besides the model incorporating $P_{active}$, we also inferred the unsteady-state *hb* kinetics using the standard two-state model (with permanently activatable gene loci) as a control (**Supplementary Fig. 5d**). By comparing the maximum likelihoods of the two models, we found that the experimental data were better fitted by the model with $P_{active}$ (**Supplementary Fig. 5e**).

#### 1.6. Modeling Bcd regulation of *hb* kinetics

Previous studies have shown that bursty *hb* transcription kinetics (during the activatable period) in early development is driven by Bcd binding on *hb* regulatory sequences[2,6]. To quantitatively understand how Bcd binding modulates *hb* transcription, we constructed a simple model to explain the time-resolved experimental data. This model describes cooperative Bcd binding at individual *hb* loci as a simple two-state kinetics, where all binding sites on the regulatory sequences are either fully occupied or fully unoccupied. The transitions between these two states (binding and unbinding) are modeled as Poisson processes, with the binding rate dependent on Bcd concentration ($C_{\text{Bcd}}$) via a power law, while the unbinding rate remains constant[14] . The probability of Bcd binding at *hb* is given by:

$$\dot{P}_{\text{bound}} = k_+ C_{\text{Bcd}}^n (1 - P_{\text{bound}}) - k_- P_{\text{bound}} \tag{S15}$$

where $n$ is the number of binding sites, $k_+$ and $k_-$ are rate constants for binding and unbinding, respectively. Assuming the binding sites are initially empty and the Bcd concentration remains constant, the Bcd binding probability evolves over time as:

$$P_{\text{bound}}(t) = \frac{C_{\text{Bcd}}^n}{C_{\text{Bcd}}^n + k_- / k_+} \left(1 - e^{-(k_+ C_{\text{Bcd}}^n + k_-)t}\right) \tag{S16}$$

When $t \to \infty$, **Equation (S16)** reaches a steady state at $P_{\text{bound}} = \dfrac{C_{\text{Bcd}}^n}{C_{\text{Bcd}}^n + k_- / k_+}$.

Since Bcd binding is a key step in triggering *hb* activation, $P_{\text{bound}}$ should be proportional to the activation rate. However, as gene activation involves additional molecular events before or after Bcd binding[2,6,8], the overall gene activation rate is linked to the Bcd binding probability by:

$$k_{\mathrm{ON}}^{-1} = aP_{\mathrm{bound}}^{-1} + \tau_0 \qquad \text{(S17)}$$

where $a$ is a proportionality constant and $\tau_0$ represents the time scale of Bcd-independent molecular events. With Bcd binding at steady state, **Equation (S17)** simplifies to:

$$k_{\mathrm{ON}} = \frac{C_{\mathrm{Bcd}}^{n}}{(a+\tau_0)C_{\mathrm{Bcd}}^{n} + ak_{-}/k_{+}} \qquad \text{(S18)}$$

This equation fitted the observed steady-state relationship between $k_{\mathrm{ON}}$ and Bcd concentration (**Fig. 5h**), from which we estimated $n \cong 5$, $a$ = 9.6 s, $\tau_0$ = 0.048 s, and $k_+/k_-$ = 6.7 × $10^{-5}$ $\mathrm{nM}^{-5}$.

**Equation (S17)** also captures the unsteady-state $k_{\mathrm{ON}}$ at the onset of each nuclear cycle. To demonstrate this, we focused on the later part of the rising phase of $k_{\mathrm{ON}}$, at which point Bcd concentration had just stabilized. In nuclear position bins with low Bcd concentration (e.g., $C_{\mathrm{Bcd}}$ = 7.5 nM at 0.45 EL), combining **Equations (S16)** and **(S17)** yields:

$$k_{\mathrm{ON}} \approx \frac{k_{+}C_{\mathrm{Bcd}}^{n}}{ak_{-}}\left(1 - e^{-(k_{+}C_{\mathrm{Bcd}}^{n}+k_{-})t}\right) \qquad \text{(S19)}$$

which fits the observed $k_{\mathrm{ON}}$ dynamics (**Fig. 5i**) and suggests that $k_{+}C_{\mathrm{Bcd}}^{n} + k_{-}$ = 3.8 $\mathrm{min}^{-1}$. Together with the estimated $k_+/k_-$, we directly determined the Bcd binding and unbinding rate constants as $k_+$ = 1.1 × $10^{-4}$ $\mathrm{min}^{-1}\mathrm{nM}^{-5}$ and $k_-$ = 1.6 $\mathrm{min}^{-1}$.

Previously, TF binding kinetics could only be measured through single-molecule tracking, which typically indicates a TF binding/unbinding timescale on the order of several seconds to a few tens of seconds[15-18]. Specifically, the genome-wide average residence time for Bcd has been reported to be relatively short, around two seconds[15,18]. However, a detailed characterization of functional binding events, i.e., those directly driving transcription, on specific genes has been largely missing. Here, our method enables a direct characterization of functional Bcd binding kinetics for *hb* activation, revealing a much longer residence time of ~40 s. This finding aligns with recent live-imaging studies of genetically modified TFs in *Drosophila* embryos[19], suggesting that functional Bcd binding events may be rare but more stable within the genome.

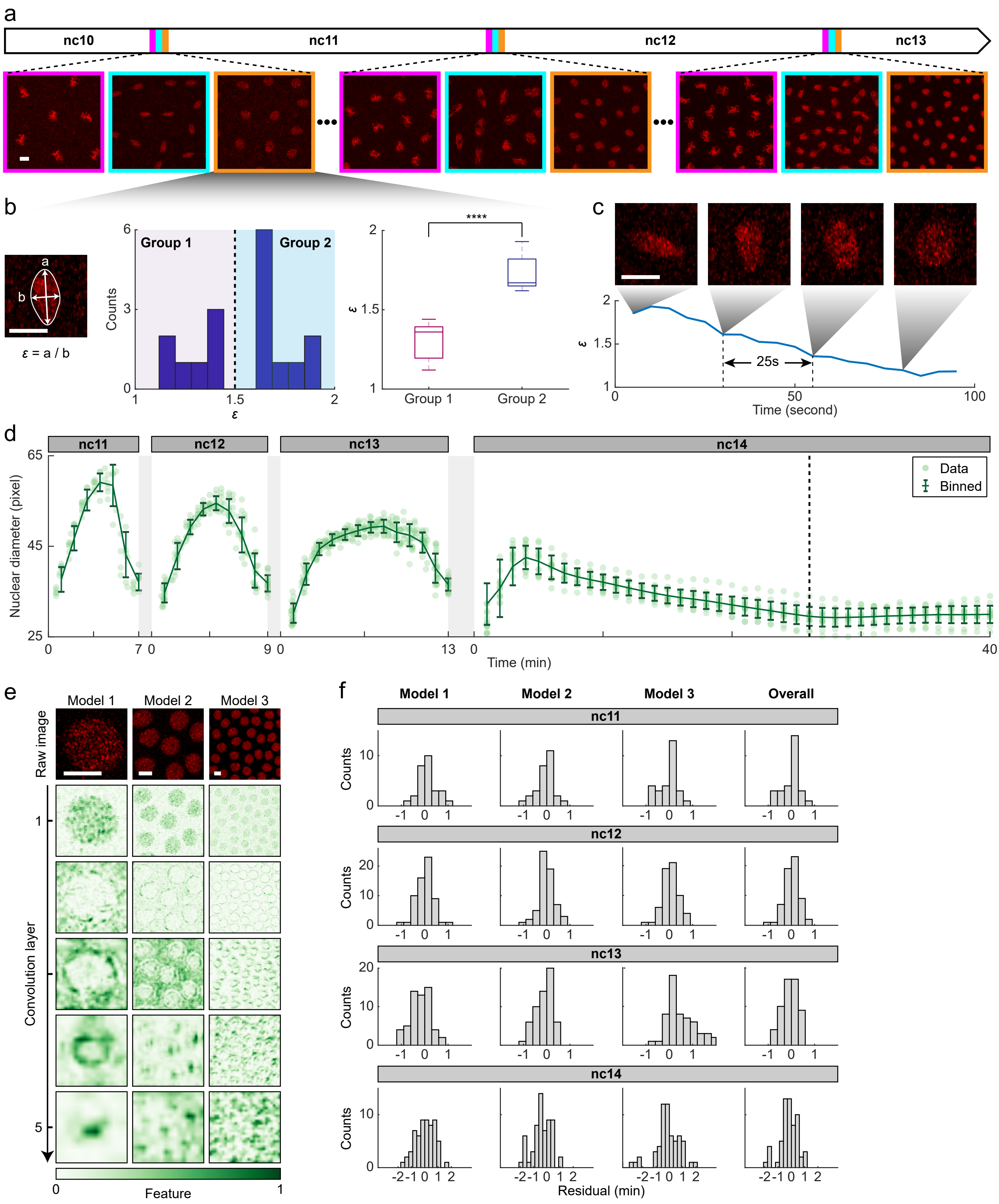


**Supplementary Fig. 1. Training and testing the histone-based time predictor. a,** Schematic for determining nuclear cycle timing by identifying nuclear division events (magenta frames) in time-lapse images of a transgenic *Drosophila* embryo (*his2av-mrfp1*) during nc10-13. Scale bar, 5 μm. **b,** Histogram of the median nuclear axial ratio ($\varepsilon$) for all live-imaging embryos at the beginning of nc11. Two groups of embryos were identified from the histogram, with distinct $\varepsilon$ (right). Statistical analysis was performed using two-sided *t*-test (*****p*<0.0001). **c,** The trend of nuclear axial ratio over time during nc10-11 division, measured from high-temporal-resolution imaging (12 fpm), with representative frames shown at the top. Timings for the two embryo groups in **b** were identified to allow estimating the time offset between them. Scale bar, 5 μm. **d,** The trend of nuclear diameter over time during nc11-14. Data from individual embryos were binned along the time axis (bin size: 1 min for nc11-13, 2 min for nc14, step size: 1 min) to show mean ± SD. *n* = 11, 18, 22, and 13 embryos for nc11-14, respectively. Dashed line indicates the end of the decline at 26 min in nc14. **e,** Feature extraction from nuclear histone images. Top: representative histone images of an nc13 embryo at different spatial scales as inputs for three independent models. Bottom: intermediate feature maps from the first five convolutional layers in each model. Scale bar, 5 μm. **f,** Histograms of residuals from individual models and the overall time predictor for each nuclear cycle. *n* = 4, 6, 4, and 3 embryos for nc11-14, respectively.

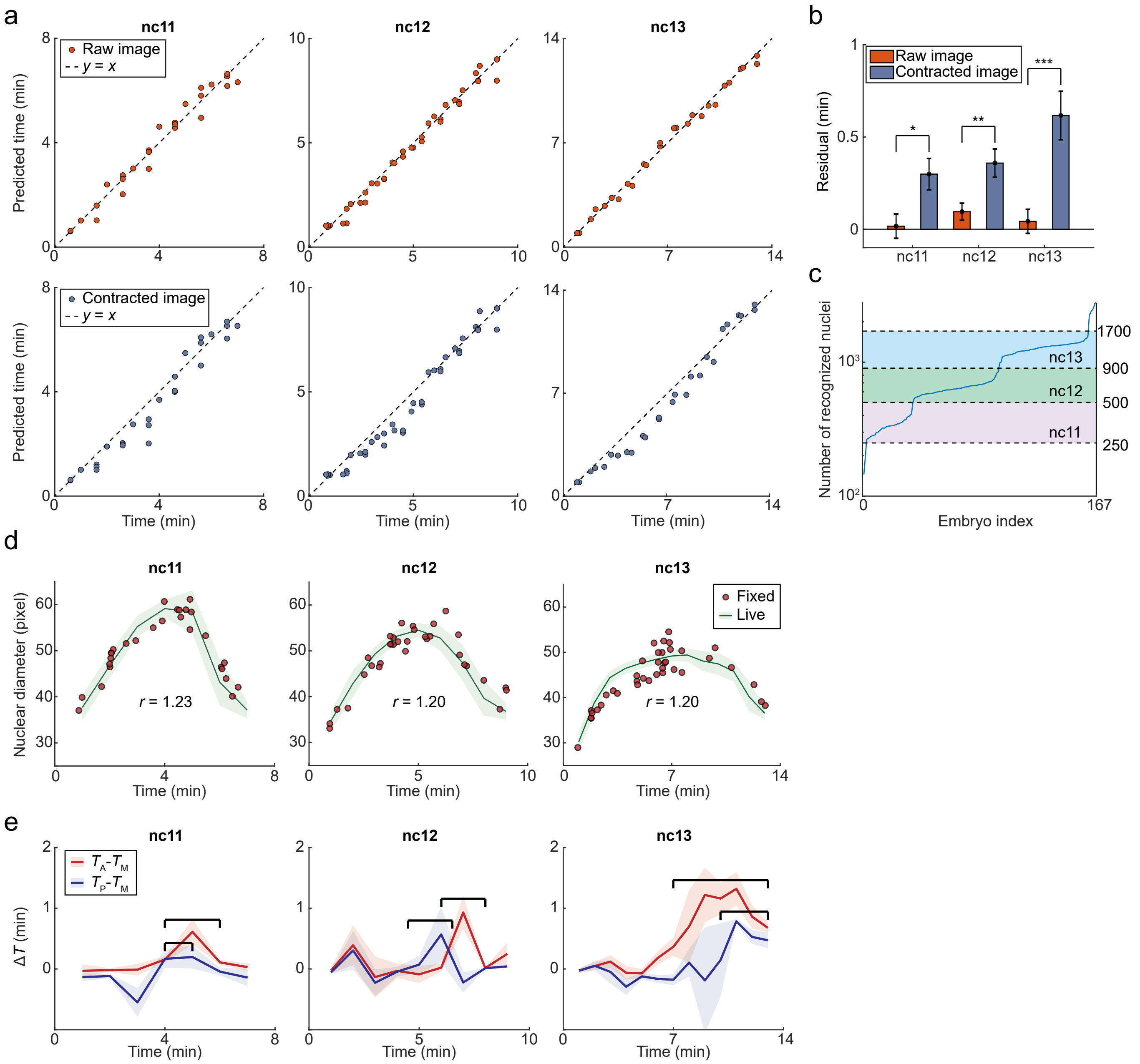


**Supplementary Fig. 2. Adapting the histone-based time inference to fixed embryos. a,** Evaluation of the impact of nuclear size shrinkage on time inference accuracy during nc11-13. For each time point in each live-imaging embryo, predicted times from raw images (top) and artificially contracted ones (contraction ratio: 0.9, bottom) were compared to ground truth, respectively. *n* = 4, 4, and 2 embryos for nc11-13, respectively. **b,** Comparison of residuals (mean ± s.e.m.) between predictions from raw and artificially contracted images for each nuclear cycle, with two-sided *t*-test (*$p<0.05$; **$p<0.01$; ***$p<0.001$). Nuclear size shrinkage significantly reduces the accuracy of time prediction. **c,** Number of recognized nuclei per embryo for 167 nc10-14 embryos, sorted in ascending order. Dashed lines indicate boundaries between nuclear cycles. **d,** Matching trends of nuclear diameter over time between live (green lines, mean ± SD) and fixed (red dots, single-embryo values) data with optimal rescaling magnitude for each nuclear cycle. Live-imaging data: *n* = 11, 18, and 22 embryos for nc11-13, respectively. Fixed-imaging data: *n* = 28, 33, and 38 embryos for nc11-13, respectively. **e,** Differences in inferred times between anterior and medial regions (red line), and between posterior and medial regions (blue line), as functions of time for each nuclear cycle. Data from multiple fixed embryos were binned along the time axis (bin size: 1 min). Shadings indicate s.e.m. Marked regions, time periods in late mitotic interphase with maximum differences in inferred times (anterior-medial difference: 4-6 min, 6-8 min, and 7-13 min for nc11-13, respectively; posterior-medial difference: 4-5 min, 4.5-6.5 min, and 10-13 min for nc11-13, respectively). *n* = 40, 41, and 49 embryos for nc11-13, respectively.

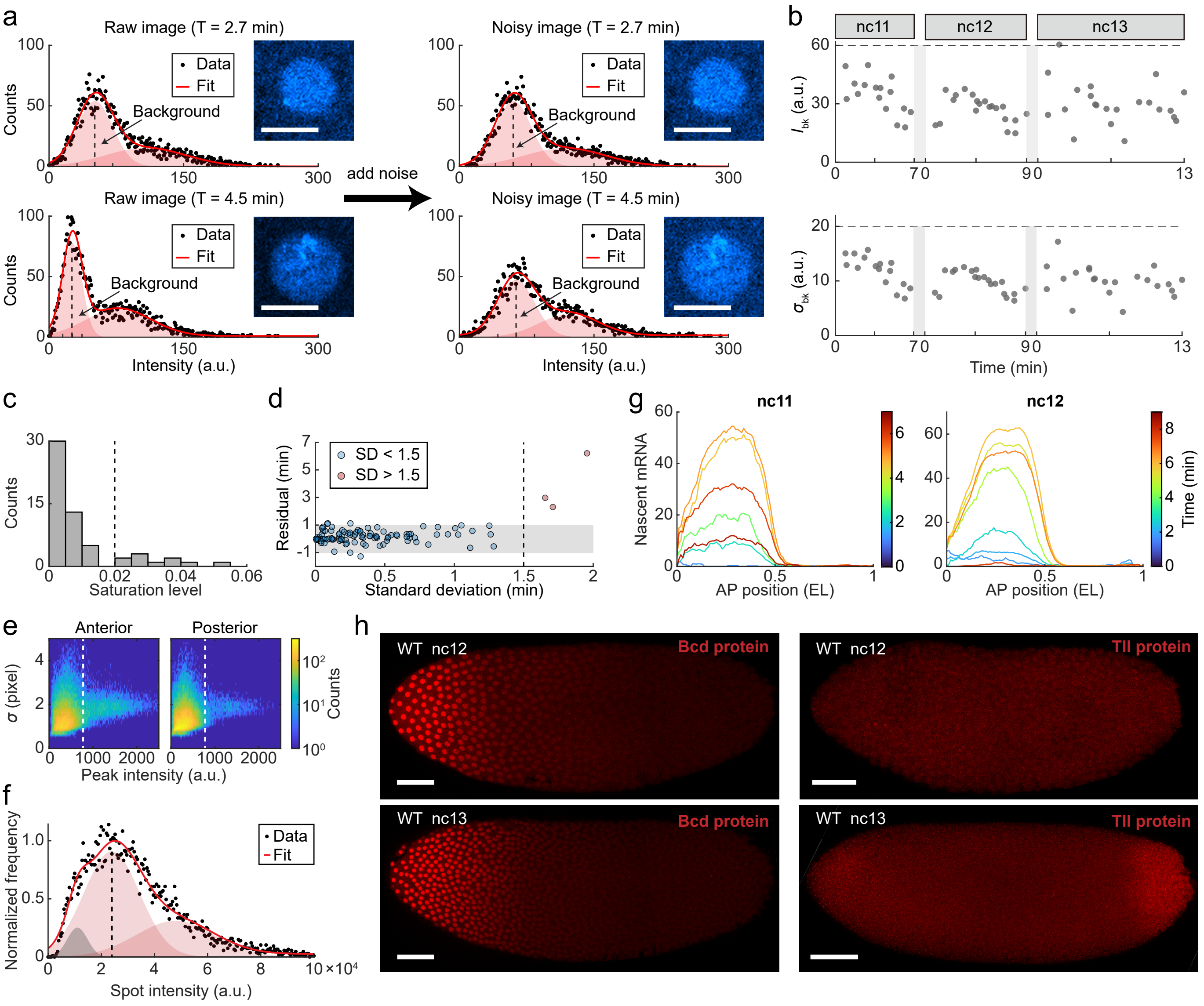


**Supplementary Fig. 3. Training and testing the DNA-based time predictor for fixed WT embryos. a,** Pixel intensity histograms from two cropped DNA images (upper and lower) before (left) and after (right) adding Gaussian noises for background calibration. The peak intensity ($I_{bk}$) and width ($\sigma_{bk}$) of background fluorescence were extracted by fitting the histogram to a sum of Gaussian functions. Insets, images before and after adding noise. Scale bars, 5 μm. **b,** Median values of $I_{bk}$ and $\sigma_{bk}$ across multiple cropped images from each training embryo in nc11-13. Upper limits for $I_{bk}$ and $\sigma_{bk}$ (dashed lines) were used to determine the appropriate Gaussian noise for each cropped image. **c,** Histogram of the average saturation level ($r_{sat}$) for different training embryos. Dashed line represents a 2% threshold used to distinguish two groups of embryos with distinct $r_{sat}$. **b-c,** $n$ = 16, 20, and 22 embryos for nc11-13, respectively. **d,** Validation of the DNA-based time predictor. For each of the 102 embryos in the test group, the residual of the final prediction is plotted against the standard deviation (SD) of nine independent predictions from three models for the anterior, medial, and posterior regions of the embryo, respectively. Embryos are divided into two groups based on an SD threshold (dashed line). Shading indicates a 1-minute accuracy range. **e,** Joint distribution of peak height and radius for candidate smFISH spots in the anterior (left) and posterior (right) regions of an embryo labeled for *hb* mRNA (>110,000 spots from a WT embryo at nc12). A threshold (dashed line) was applied to distinguish real *hb* mRNA spots from false-positive particles. **f,** Intensity histogram of *hb* smFISH spots in the anterior region of the embryo (>50,000 spots). The intensity corresponding to a single mRNA molecule was identified by fitting the histogram to a sum of Gaussian functions. **g,** Spatial profiles of endogenous *hb* transcription over time during nc11 and nc12, binned along the AP axis (bin size: 0.06 EL, step size: 0.01 EL) and time (bin size: 2 min, step size: 1 min). **h,** Confocal images of WT embryos labeled for Bcd protein (left) and Tll protein (right) at nc12 and nc13, respectively. Scale bars, 50 μm. Compared to maternal Bcd, which exhibits a strong signal during nc12-13, the Tll signal is present in nc13 but not in nc12.

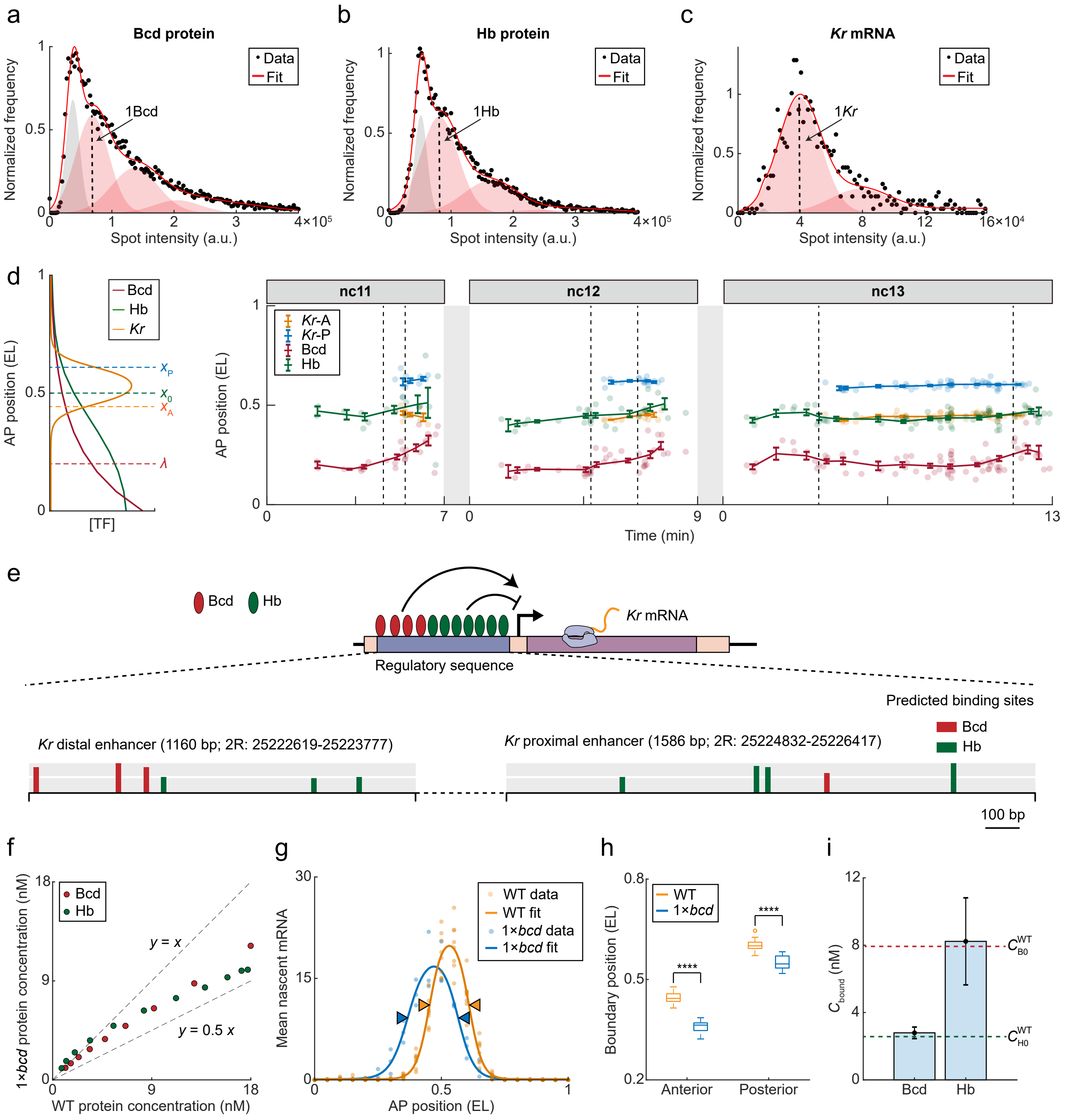


**Supplementary Fig. 4. Quantitative analysis of dynamic *Kr* regulation using fixed embryos. a-b,** Intensity histograms of cytoplasmic Bcd (**a**) and Hb (**b**) spots in the anterior region of the embryo (each >20,000 spots from a WT embryo at nc13). The intensities corresponding to individual Bcd and Hb protein molecules were identified by fitting each histogram to a sum of Gaussian functions, respectively. **c,** Intensity histogram of *Kr* smFISH spots in the medial region of the embryo (>20,000 spots from a WT embryo at nc13). The intensity corresponding to a single mRNA molecule was identified by fitting the histogram to a sum of Gaussian functions. **d,** Anterior and posterior *Kr* expression boundaries ($x_A$ and $x_P$), decay length of the Bcd profile ($\lambda$), and boundary position of the Hb profile ($x_0$) as functions of time. Data were binned along the time axis (bin size: 2 min, step size: 1 min) to show mean ± s.e.m. Dashed lines mark *Kr* expression periods ($k > 0$) in each nuclear cycle. Bcd analysis: $n$ = 25, 36, and 76 embryos for nc11-13, respectively; Hb analysis: $n$ = 17, 29, and 60 embryos for nc11-13, respectively; *Kr* analysis: $n$ = 11, 17, and 57 embryos for nc11-13, respectively. **e,** Strong Bcd and Hb binding sites (log-odds score > 5) on distal and proximal enhancers of the *Kr* gene, predicted using a PWM motif analysis. Data were plotted using inSite (https://www.cs.utah.edu/~miriah/insite/), with colors representing binding site identity and bar height denoting the log-odds of binding. **f,** Comparison of average Bcd and Hb concentrations at different AP positions (0.2-0.7 EL) between WT ($n$ = 53) and 1×*bcd* ($n$ = 11) embryos during the activatable period (3.8-11.5 min, $k > 0$) in nc13. **g,** Average nuclear *Kr* transcription as a function of the AP position for WT ($n$ = 8) and 1×*bcd* ($n$ = 3) embryos during 9.5-10.5 min in nc13. Data were fitted to a product of two logistic functions to extract expression boundaries (arrows). **h,** Comparison of anterior and posterior *Kr* expression boundaries between WT ($n$ = 50) and 1×*bcd* ($n$ = 7) embryos during the activatable period (3.8-11.5 min, $k > 0$) in nc13. Two-sided *t*-test was applied between strains (****$p < 0.0001$). **i,** Average Bcd and Hb concentrations at *Kr* boundaries for 1×*bcd* embryos ($n$ = 11) during the activatable period (3.8-11.5 min, $k > 0$) in nc13 (mean ± s.e.m.). Average Bcd and Hb concentration thresholds for WT nc13 embryos from **Fig. 4h** are shown as reference.

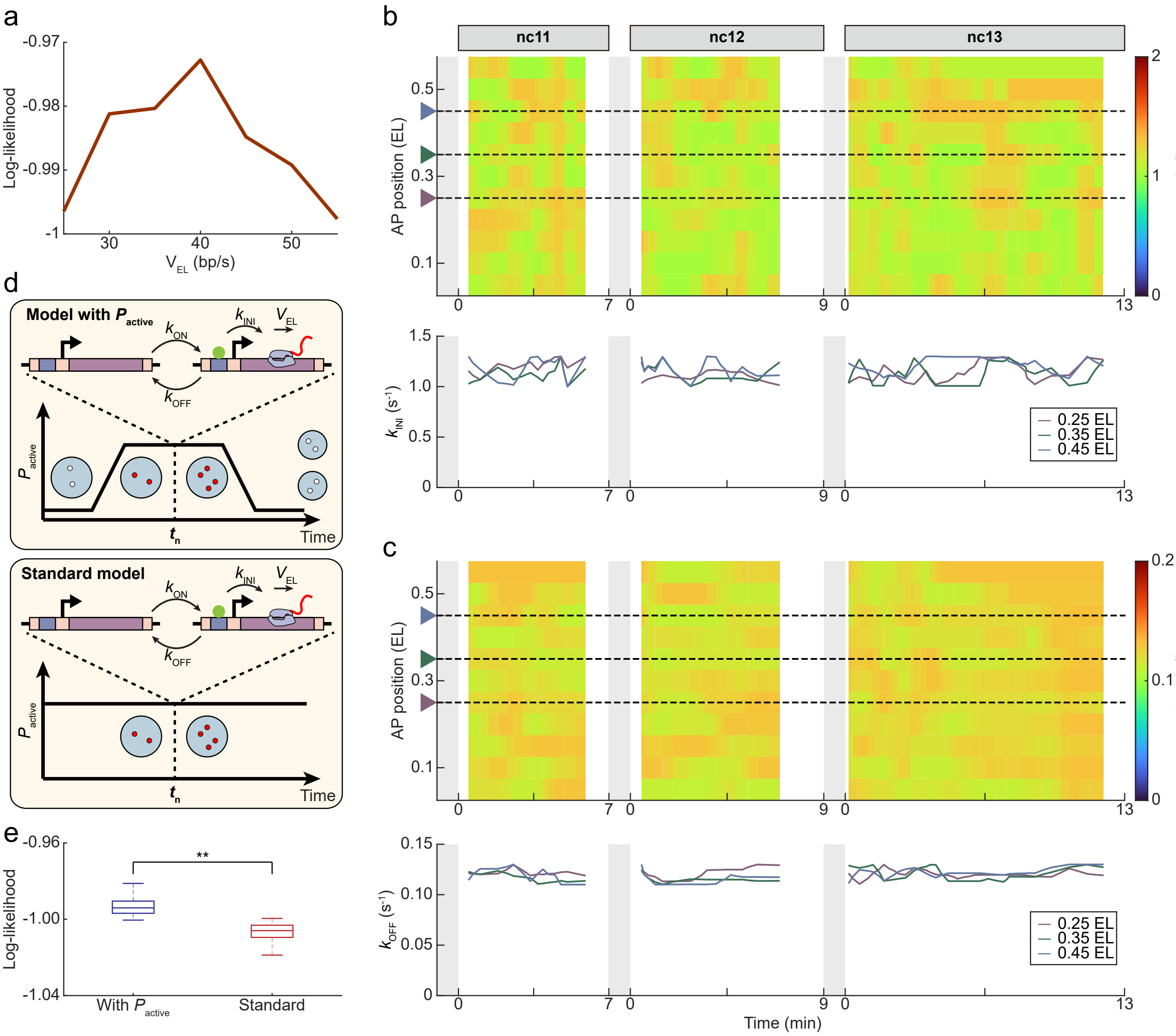


**Supplementary Fig. 5. Estimating kinetic parameters of the unsteady-state *hb* transcription model. a,** Fitting likelihood as a function of elongation speed. Fitting was applied to data from the anterior *hb* expression region (0.2-0.4 EL) during nc13. **b-c,** Spatiotemporal profiles of transcription initiation rate (**b**) and gene inactivation rate (**c**) across nc11-13 for the AP position range of 0.05-0.55 EL. Data from three specific AP positions (0.25, 0.35, and 0.45 EL, marked by dashed lines) are individually plotted as functions of time (bottom). **d,** Schematics of our two-state model with time-varying $P_{active}$ compared to the standard two-state model with $P_{active} \equiv 1$. **e,** Likelihood comparison between the standard two-state model and our two-state model with time-varying $P_{active}$. Fittings were applied to data from the anterior *hb* expression region (0.2-0.4 EL) during nc13. Two-sided *t*-test was applied between models (**$p < 0.01$).

**Supplementary Table 1. Configuration of CNN models.**

<table>
<tr><th colspan="5">CNN model Configuration</th></tr>
<tr><td>A</td><td>B</td><td>C</td><td>D</td><td>E</td></tr>
<tr><td>conv5-32</td><td>conv5-32</td><td>conv5-32</td><td>conv5-32</td><td>conv5-32</td></tr>
<tr><td colspan="5">maxpool</td></tr>
<tr><td>conv5-64</td><td>conv5-64</td><td>conv5-64</td><td>conv5-64</td><td>conv5-64</td></tr>
<tr><td colspan="5">maxpool</td></tr>
<tr><td>conv5-128</td><td>conv5-128</td><td>conv5-128</td><td>conv5-128</td><td>conv5-128</td></tr>
<tr><td colspan="5">maxpool</td></tr>
<tr><td rowspan="6">None</td><td>conv5-256</td><td>conv5-256</td><td>conv5-256</td><td>conv5-256</td></tr>
<tr><td colspan="4">maxpool</td></tr>
<tr><td rowspan="4">None</td><td>conv5-128</td><td>conv5-128</td><td>conv5-512</td></tr>
<tr><td colspan="3">maxpool</td></tr>
<tr><td rowspan="2">None</td><td>conv5-64</td><td>conv5-256</td></tr>
<tr><td colspan="2">maxpool</td></tr>
<tr><td colspan="5">BN</td></tr>
<tr><td colspan="5">FC-500</td></tr>
<tr><td colspan="5">FC-128</td></tr>
<tr><td colspan="5">FC-64</td></tr>
<tr><td colspan="5">FC-32</td></tr>
<tr><td colspan="5">FC-16</td></tr>
<tr><td colspan="5">FC-1 (linear)</td></tr>
</table>

<table>
<tr><th rowspan="2">CNN models 1-3</th><th colspan="3">nc11</th><th colspan="3">nc12</th><th colspan="3">nc13</th><th colspan="3">nc14</th></tr>
<tr><th>1</th><th>2</th><th>3</th><th>1</th><th>2</th><th>3</th><th>1</th><th>2</th><th>3</th><th>1</th><th>2</th><th>3</th></tr>
<tr><td>Histone-based</td><td>A</td><td>A</td><td>A</td><td>C</td><td>C</td><td>D</td><td>C</td><td>C</td><td>E</td><td>C</td><td>C</td><td>B</td></tr>
<tr><td>DNA-based</td><td>C</td><td>A</td><td>C</td><td>C</td><td>C</td><td>C</td><td>C</td><td>C</td><td>C</td><td colspan="3">None</td></tr>
</table>

* Five configurations (A-E, shown in different columns from left to right) were used to train Histone- and DNA-based CNN models for nc11-14, respectively. The parameters for the convolutional layers are denoted as “conv <kernel size> - <number of kernels>”.
